# Surface Wetting Is a Key Determinant of α-Synuclein Condensate Maturation

**DOI:** 10.1101/2024.06.06.597302

**Authors:** Rebecca J. Thrush, Devkee M. Vadukul, Siân C. Allerton, Marko Storch, Francesco A. Aprile

## Abstract

α-Synuclein self-assembles into amyloid fibrils during neurodegeneration. The protein can also self-assemble via liquid-liquid phase separation to form biomolecular condensates. The link between these processes is evident, as α-synuclein condensates can mature into amyloids. However, the mechanisms driving this maturation remain largely unknown, particularly when incorporating pathological post-translational modifications known to affect α-synuclein self-assembly in the absence of LLPS, such as N-terminal truncation. Moreover, condensates are primarily studied as isolated entities; however, it is increasingly evident that they interact with various cellular components and surfaces. Here, we developed a microscopy-based quantitative real-time imaging protocol to investigate how N-terminal truncation influences α-synuclein condensate formation, well surface wetting, and maturation. We found that increasing α-synuclein truncation, which reduces N-terminal hydrophobicity, inhibits condensate sedimentation, enhances surface wettability, and accelerates maturation. Additionally, by increasing well surface hydrophobicity we decreased α-synuclein condensate wettability, delaying their maturation. Thus, we propose that enhanced wettability, which increases the condensate surface-to-volume ratio, promotes α-synuclein nucleation at the condensate-bulk solution interface, thereby accelerating maturation. Our results reveal distinct mechanistic roles for α-synuclein N-terminal residues and indicate that condensate wetting on cellular surfaces, such as synaptic vesicles, may drive toxic aggregate formation during neurodegeneration.

## Introduction

The intrinsically disordered protein (IDP) α-synuclein (α-syn) exists as a monomer and multimers in neurons, where it mediates neurotransmitter release and synaptic function.^1-6^ α-Syn is also found as insoluble amyloid fibrils with a characteristic cross β-sheet rich core structure.^7,8^ The aberrant accumulation of α-syn amyloid fibrils is the major hallmark of neurodegenerative diseases including Parkinson’s disease (PD), dementia with Lewy bodies, and multiple system atrophy.^9,10^

Many post-translational modifications (PTMs) affect α-syn amyloid aggregation and PD pathology.^11-20^ These PTMs include acetylation, phosphorylation, and terminal domain truncation.^12-15^ One PTM that remains largely uncharacterized is N-terminal truncation, which has been observed both in vitro^21,22^ and in vivo.^13-15,23-26^ The N-terminus of α-syn plays key roles in both the physiological and pathological functions of the protein, e.g., the domain acts as a membrane anchor region,^27,28^ modulating synaptic vesicle fusion to the presynaptic terminal.^4,29^ Yet, in vitro evidence shows that vesicle bound α-syn is often predisposed to self-association, and can nucleate free monomer to further promote aggregation.^27,30^ The N-terminus also plays a role in the growth phase of amyloid aggregation, specifically elongation and secondary nucleation.^31,32^ Thus, unsurprisingly, N-terminal truncations and deletions have been shown to alter α-syn aggregation kinetics, amyloid morphology and stability, membrane binding, and toxicity, relative to the full-length (FL) protein.^33-38^

Emerging evidence demonstrates that α-syn can undergo liquid-liquid phase separation (LLPS), resulting in the formation of biomolecular condensates.^39-45^ At the early stages of maturation, the condensates exhibit liquid-like properties, allowing α-Syn to diffuse within them and exchange with the surrounding environment.^39,44^ These liquid-like properties are likely facilitated by weak multivalent interactions between α-syn molecules.^46,47^ The condensates can also grow via fusion events (coalescence) and Ostwald ripening.^39,44^ With time, the condensates can mature into a solid-like gel, where composing α-syn molecules exhibit reduced mobility.^39,44^ This liquid-to-solid transition ultimately leads to the formation of amyloids.^39,43,44^ Several environmental conditions can promote α-syn LLPS. A molecular crowding agent, commonly poly-ethylene glycol (PEG), is often necessary, although evaporation, poly-cations, metal ions or other amyloidogenic proteins, e.g., tau, can also induce LLPS.^39-44^ Interestingly, recent evidence indicates that, under certain conditions, the condensate-bulk solution interface can act as a catalyst for the aggregation of amyloidogenic proteins, including α-syn.^45,48,49^

The exact protein regions and interactions regulating α-syn LLPS remain largely unknown. However, the critical protein concentration required for LLPS is significantly reduced by lowering pH, increasing salt, or introducing metal ions.^39,40^ Similarly, interaction of the negative C-terminus of α-syn with positively charged regions of tau or poly-L-lysine (PLK) induces complex coacervation; the co-LLPS of two opposingly charged molecules.^42^ Moreover, a variant possessing only the core residues 30–110 and C-terminally truncated α-syn both undergo enhanced LLPS and aggregation, relative to FL α-syn.^39,41^ Together, these findings indicate that electrostatic interactions mediate α-syn condensate formation, suggesting a key role for the terminal domains. Furthermore, b-syn, which lacks the hydrophobic eight residue stretch located in the non-amyloid-b component (NAC) domain of α-syn, does not phase separate or aggregate, indicating that this domain is critical for LLPS.^39^Amyloid aggregation is significantly affected by conditions that promote LLPS. For several proteins, including α-syn, tau and heterogeneous nuclear ribonucleoprotein A1 (hnRNPA1), amyloid aggregation is accelerated under LLPS conditions.^39,46,48,50,51^ This effect is likely due to increased local protein concentration and enhanced nucleation. Furthermore, various disease factors known to promote α-syn aggregation in the absence of LLPS (*e*.*g*., Cu^2+^ and Fe^3+^) also accelerate the onset of LLPS and the subsequent liquid-to-solid transition.^39^ However, conditions that promote LLPS can also inhibit amyloid aggregation, as observed for the 42-residue-long amyloid-b peptide, which can be sequestered and stabilized within condensates formed from the low complexity domains of the DEAD-box proteins.^52^

The ability of biomolecular condensates to wet surfaces is believed to be a biologically relevant mechanism.^53-55^ In cells, condensate wetting on biological surfaces, such as lipid membranes, can affect their liquid-to-solid transition, potentially contributing to amyloid formation and cellular toxicity.^53,55^

Despite the importance of the N-terminus, molecular details of how specific N-terminal regions regulate the nucleation and propagation mechanisms of α-syn self-assembly remain unclear. In this study, we used a microscopy-based approach to identify sequence determinants within the N-terminus that regulate the balance between the surface wettability and maturation of α-syn condensates.

## Results

### Design of the N-terminally Truncated α-Syn Variants

Several N-terminal truncations have been associated with PD. We selected three different truncations (Table S2): 5-140, isolated from PD brains and following *in vitro* incubation,^13,22^ 11-140, found in Lund human mesencephalic cells,^15^ and 19-140, found in human appendices.^14^ The N-terminal region of these variants differs from the FL protein in both charge and hydrophobicity. 5-140 α-syn has a relative charge (ΔCharge) of +1, while both 11-140 and 19-140 α-syn have a ΔCharge of -1, relative to FL α-syn. In terms of N-terminal hydrophobicity, FL α-syn is the most hydrophobic, followed by 11-140 and then 5-140 α-syn. 19-140 α-syn has the lowest hydrophobicity. We recombinantly expressed and purified all three variants, alongside FL α-syn, as confirmed by ESI-MS (Supplementary Fig. 1). Using far-UV CD spectroscopy we found that, similar to FL α-syn, all truncated variants consist primarily of random coil secondary structure (Supplementary Fig. 2).

### N-terminal truncation does not affect the ability of α-syn to undergo LLPS

α-Syn has been reported to naturally interact with basic proteins, such as synapsins.^42,56^ In vitro, α-Syn can undergo LLPS when incubated with the polycation PLK and a molecular crowding agent, *e*.*g*., PEG, mimicking physiological conditions.^42^ To investigate how N-terminal truncation affects α-syn LLPS, we generated phase diagrams for our α-syn variants. We incubated varying concentrations of each protein with 10 % PEG and different concentrations of PLK for 15 min, when the highest number of condensates was observed (Fig. 1a,b). Then, we imaged the bulk solution of each sample using differential interference contrast (DIC) microscopy, designed a General Analysis 3 (GA3) recipe that estimates condensate number and area (Supplementary Fig. 3), and plotted the number of condensates in solution as a function of both α-syn and PLK concentration (Fig. 1a and Supplementary Fig.4). These values were combined over 3 z-stack images acquired in solution per sample (Fig. 1a). The distance between the planes (200 mm) was chosen to be ~ 100-fold greater than the average condensate diameter (~ 2.5 µm) to ensure no condensate is captured in more than one image.

**Figure 1.**
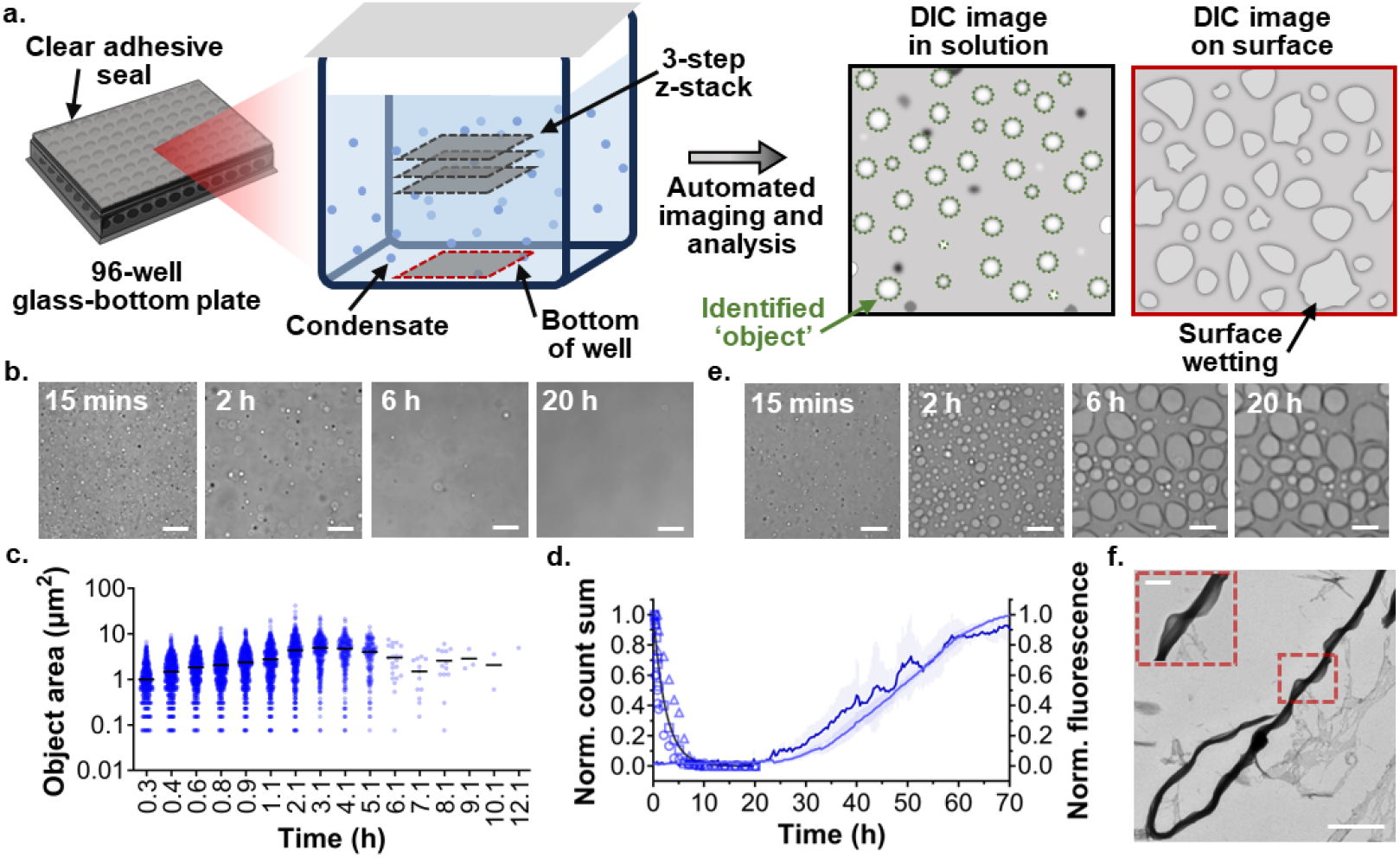
Condensate and amyloid formation of FL α-syn under LLPS-promoting conditions. **(a)** A diagram outlining the methodology used to monitor α-syn liquid condensate formation, sedimentation and maturation. Samples, loaded in a 96-well glass-bottom plate sealed with a clear film, are incubated at 37°C. Automated 3-step z-stack imaging in solution, alongside bottom of well imaging, is performed at selected time points over 20 h. Solution image analysis is then performed to quantify the number and area of all in focus objects within the image. **(b)** Representative DIC images acquired in solution during FL α-syn incubation. Scale bars represent 25 μm. **(c)** DIC microscopy-derived object area distribution over time. Each timepoint was compiled over three z-stack images. Mean areas are indicated by a solid black line. **(d)** Normalized total object count (left Y-axis) and normalized ThT intensity (right Y-axis) as a function of time. Three biological repeats are shown for the normalized total object count data (circles, squares, or triangles), which are globally fitted using a one-phase decay model (black line). Two biological repeats are shown for the normalized ThT aggregation data (dark and light blue lines), where each biological repeat is the mean of three technical replicates and error bars represent the standard deviation of the mean. **(e)** Representative DIC images acquired at the bottom of the well during FL α-syn incubation. Scale bars represent 25 μm. **(f)** Representative TEM image, with magnification, of an aliquot collected at the endpoint of a LLPS ThT aggregation assay. Scale bars represent 2 μm and 500 nm, respectively.

FL α-syn formed condensates at either 40 or 60 mM with 25 mM PLK, and at 60 mM with 50 mM PLK (Supplementary Fig. 4 and 5). To verify that condensates contained FL α-syn, we performed fluorescence microscopy on a sample of 60 mM FL α-syn, 1 % of which was rhodamine labelled, and 25 mM PLK. Fluorescence emission was specifically localized within the condensates, confirming the presence of α-syn (Supplementary Fig. 6). We repeated this analysis on 5-140, 11-140 and 19-140 α-syn. We found that all truncated variants formed condensates, with a similar dependency on the ratio of protein:PLK as FL α-syn (Supplementary Fig. 4 and 7─9). This suggests that residues 1–18 are not required for condensate formation, consistent with existing evidence that this complex coacervation mechanism requires interaction between PLK and the C-terminus of α-syn and that residues 30–110 are sufficient for α-syn LLPS.^39,42^

### N-terminal truncation accelerates α-syn amyloid formation under LLPS conditions

We next investigated whether N-terminal truncation affects the time-evolution of α-syn condensates. We prepared solutions containing 60 mM α-syn, 10 % PEG, and 25 mM PLK, as we observed significant condensate formation under this condition for all α-syn variants (Supplementary Fig. 4). We monitored condensate maturation in the bulk of the solution using DIC microscopy and then quantified condensate area and number variation with time (Fig. 1a). Additionally, we imaged the bottom surface of the well at each time point to monitor any condensate or aggregate sedimentation during incubation (Fig. 1a).

First, we examined FL α-syn and found that condensates increased in size (from ~ 1 to ~ 5 μm^2^ area) during the first ~ 3 h of incubation (Fig. 1b,c and Supplementary Fig. 10a). This increase in size was attributed in part to coalescence, which was detected after ~ 20 min, and also evidences the liquid-like nature of early α-syn condensates (Supplementary Fig. 11). After ~ 3 h, the condensates stopped growing (Fig. 1b,c and Supplementary Fig. 10a), possibly because their low number reduced the likelihood of proximity-dependent growth events (*i*.*e*., coalescence and Ostwald ripening). Alternatively, the condensates may have transitioned into a gel-like phase, preventing further growth.^39,44^

In contrast, the number of condensates in the bulk solution decreased continuously after 15 min (Fig. 1b,d and Supplementary Fig. 10b). By 20 h, very few condensates were visible in solution. This behavior was partially attributed to coalescence, which would decrease the number of condensates. We also imaged the bottom of well throughout incubation and found a progressive increase in the number of condensates wetting the well surface (Fig. 1e), indicating sedimentation also contributes to the decay in condensate number in solution. When manually imaging at 1.5 h we observed fusion events between condensates in solution and surface-wetted condensates (Supplementary Fig. 12), explaining the progressive increase in surface-wetted condensate size over time (Fig. 1e). We did not observe condensates, surface wetting, or aggregates in any of the control samples, i.e., 60 mM FL α-syn alone, 25 mM PLK alone or LLPS buffer alone (Supplementary Fig. 13 and 14).

To investigate the transition of FL α-syn condensates into amyloids, we performed fluorescence emission measurements in a microplate reader using the amyloid-sensitive probe Thioflavin-T (ThT). We prepared samples containing 60 mM α-syn under the same LLPS-inducing conditions discussed above. We then recorded ThT fluorescence intensity over time as a readout of amyloid formation. To mimic the microscope set-up, we took readings at the well center every 20 min with 30 flashes per well, minimizing sample agitation and overheating. Aggregation commenced after ~ 20 h (Fig. 1d). In the absence of PLK, we did not observe any significant aggregation within the time frame of this experiment (Supplementary Fig. 15a). These data are consistent with our microscopy observations (Fig. 1b,d, Supplementary Fig. 13 and 14) and existing literature evidence that LLPS can accelerate amyloid aggregation.^39,46,48,50,51^

To characterize the aggregates formed following LLPS, we analyzed the endpoint of this assay. Microscopy revealed ThT-positive aggregates, including large objects with protruding fibrils (Supplementary Fig. 16 and Supplementary Movie 1). TEM also confirmed the presence of large, twisted fibrils (Fig. 1f). To understand the composition and secondary structure of these aggregates we isolated and analyzed the insoluble protein fraction. We found that this fraction contained α-syn and displayed b-sheet secondary structure, as assessed by dot-blot and far-UV CD spectroscopy, respectively (Supplementary Fig. 17). Altogether these data indicate that amyloid aggregates are formed under LLPS conditions. In the absence of PLK, we observed a small number of thin fibrils by TEM, consistent with the ThT intensity data (Supplementary Fig. 18).

The changes in 5-140 α-syn condensate size and number over time were similar to those observed for FL α-syn (Fig. 2a and Supplementary Fig. 19–21). In contrast, 11-140 and 19-140 α-syn condensates persisted in solution for longer (Fig. 2b,c). Although their normalized growth rates were slower than FL α-syn, their maximum condensate sizes were larger (Supplementary Fig. 19b and 21), suggesting increased coalescence and Ostwald ripening. Condensate growth plateaued for all variants when ~ 30 % condensates remained in solution (Fig. 2b,c and Supplementary Fig. 19), supporting our earlier hypothesis that condensate growth plateaus when their number falls below a critical threshold. This indicates that 11-140 and 19-140 α-syn condensates reach a larger size due to their prolonged existence in solution, which may be a consequence of altered sedimentation dynamics. In fact, increasing truncation reduces N-terminal hydrophobicity (Table S2), which may enhance water retention within condensates, lowering their density and thereby slowing sedimentation.

**Figure 2.**
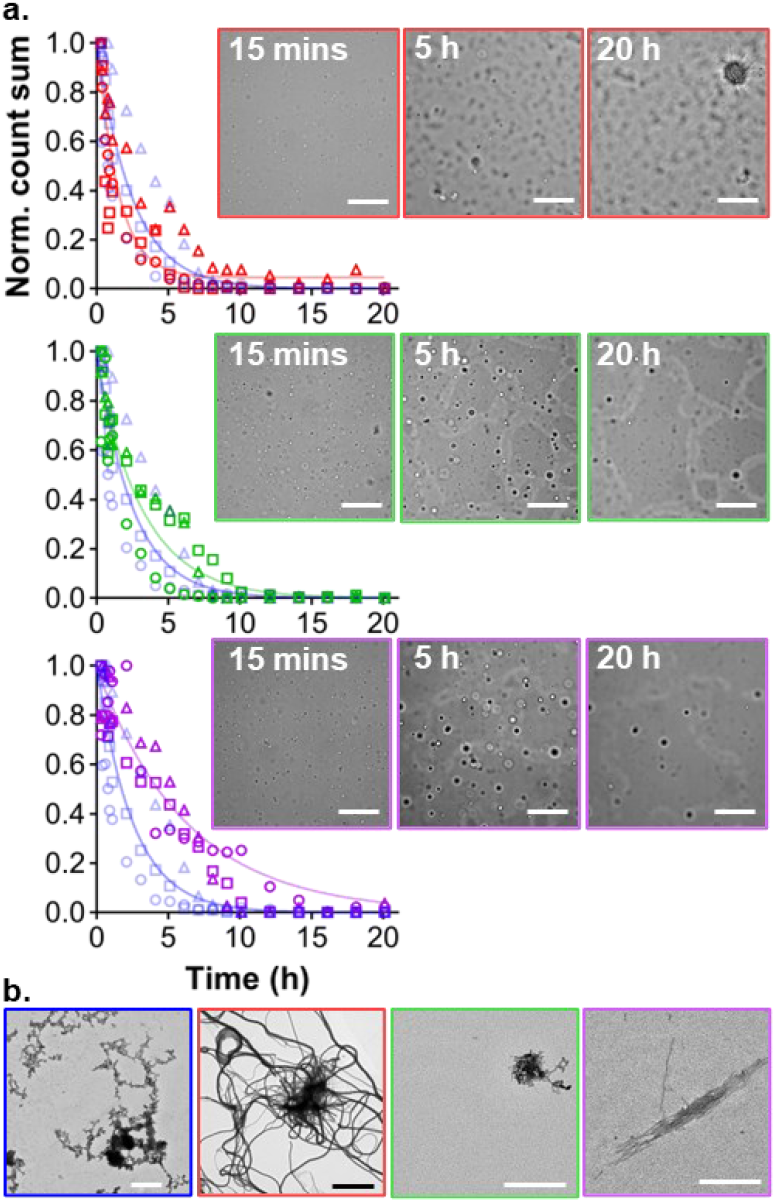
Condensate and amyloid formation of truncated α-syn under LLPS-promoting conditions. **(a)** Normalized total object count over time for 5-140 (red), 11-140 (green) and 19-140 (purple) α-syn. Three individual biological repeats are shown (circles, squares or triangles). Data are globally fitted using a one-phase decay model (solid lines). FL α-syn data (semi-transparent blue symbols and line) are shown for comparison. Overlayed are representative DIC images acquired via automation at the bottom of the well during incubation (scale bars represent 50 μm). **(b)** Representative TEM images of 5-140 (red), 11-140 (green) and 19-140 (purple) α-syn aliquots taken after 20 h incubation. Scale bars represent 400 nm, except for 5-140 α-syn whose scale bar is 4 μm.

Regarding 5-140 α-syn, the appearance of non-spherical objects in solution after ~ 2 h indicates the formation of solid aggregates (Supplementary Fig. 21). Bottom of well imaging also indicated accelerated aggregation for 5-140 α-syn as a large network of fibrils was observed at 20 h (Fig. 2a and Supplementary Fig. 22). Additionally, at the end of incubation long, thick fibril clumps could be seen by TEM (Fig. 2b).

In contrast, bottom of well microscopy of 11-140 and 19-140 α-syn showed extensive surface wetting, comparable to the FL protein (Fig. 2a). Both truncations resulted in a greater surface area of wetted condensates, which also contained small aggregates (Fig. 2a and Supplementary Fig. 22). The morphology of these condensate was less spherical, and their borders had substantially lower contrast (seen at 20 h in Supplementary Fig. 22), indicating a thinner condensate layer. This suggests condensates of 11-140 and 19-140 α-syn have a lower contact angle on the surface and thus increased wettability, relative to FL α-syn. Given that the well surface is glass and thus hydrophilic, this increased wettability is likely due to the decreased N-terminal hydrophobicity of the truncated α-syn variants.

Finally, we examined amyloid formation under LLPS-promoting conditions using the ThT intensity assay. We observed faster aggregation kinetics for all truncated variants, relative to FL α-syn (Fig. 3a and Supplementary Fig. 15b). DIC, ThT fluorescence and electron microscopy at the endpoint of aggregation revealed extensive thick fibril networks for all variants (Fig. 3b and Supplementary Movie 2─4). The insoluble protein fractions were α-syn positive and contained b-sheet secondary structure, indicative of amyloids (Supplementary Fig. 17).

**Figure 3.**
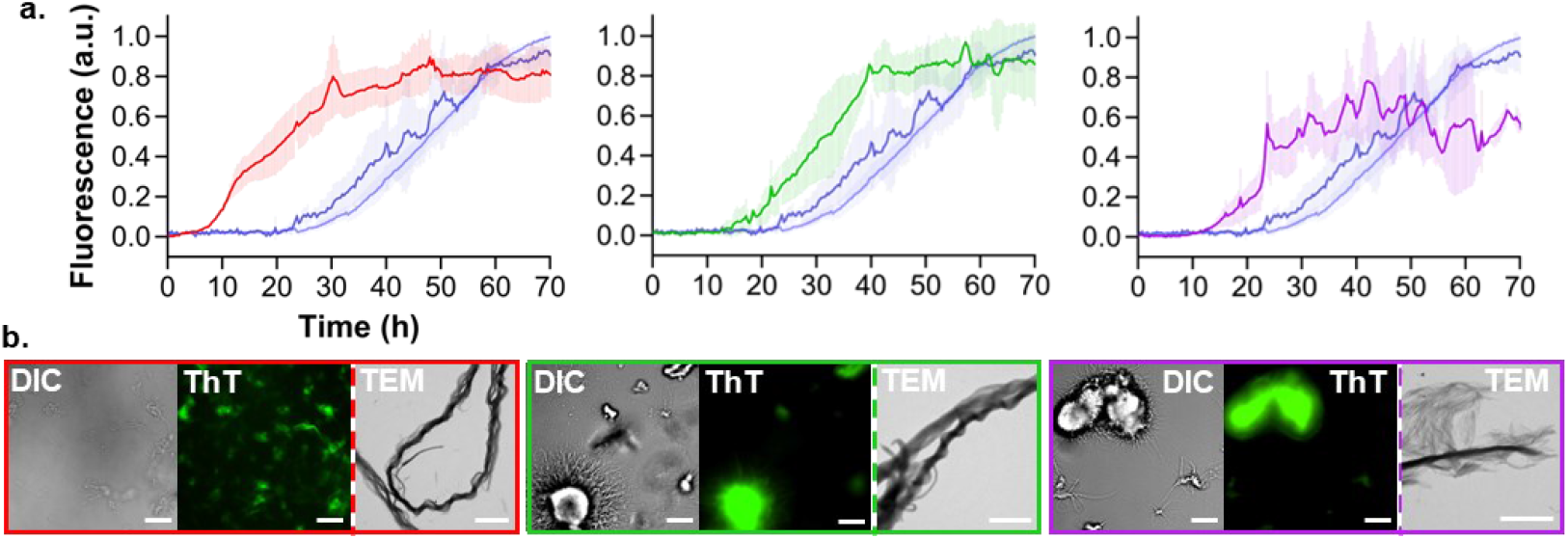
LLPS-induced amyloid formation of N-terminally truncated α-syn. **(a)** Normalized ThT intensity of 5-140 (red), 11-140 (green) or 19-140 (purple) α-syn shown alongside the FL α-syn data (semi-transparent blue) as comparison. Two (FL α-syn) or one (truncated α-syn) individual biological repeats are shown. Each repeat is the mean of three technical replicates, error bars represent the standard deviation of the mean. **(b)** Representative DIC, ThT fluorescence, and TEM images taken at the bottom of the well at the endpoint of the assay. ThT fluorescence scale bars represent 25 μm; TEM scale bars represent 3 μm.

When truncated α-syn was incubated in the absence of PLK, did we not observe condensates, surface wetting, or aggregates within 20 h by DIC microscopy (Supplementary Fig. 21 and 23). A small number of thin fibrils were formed at the end of the ThT intensity assay, consistent with delayed aggregation kinetics relative to phase-separated α-syn (Supplementary Fig. 15b and 24).

Together our data demonstrate that N-terminal truncation accelerates aggregation under conditions that induce LLPS. With regards to 11-140 and 19-140 α-syn, this may be a consequence of the increased surface wettability of their condensates. Interestingly, 5-140 α-syn aggregation initiates prior to substantial condensate sedimentation and surface wetting, indicating surface wettability is not responsible for the accelerated maturation of this variant.

### Increasing surface wettability accelerates α-syn amyloid formation

To explore the role of surface wetting in condensate maturation, we incubated FL α-syn in 8-well slides with coverslips of varying hydrophobicity. Specifically, we used slides with (i) a glass coverslip (hydrophilic surface), (ii) a polymer coverslip (mildly hydrophobic surface), and (iii) a BioInert polymer coverslip, which is treated to render the surface highly hydrophobic and inert to biomolecules. As the surface wettability of a given protein condensate is largely determined by the balance between protein-protein interactions within the condensate versus condensate-surface interactions, we hypothesized that increasing surface hydrophobicity would disfavor condensate-surface interactions, thereby decreasing wettability.

We prepared a solution of 60 µM α-syn and 25 µM PLK in LLPS buffer, supplemented with 20 µM ThT. Since direct fluorescence intensity measurements (*i*.*e*., in a microplate reader) were not possible in the well slides, we imaged the samples using fluorescence microscopy and quantified the mean intensity of each image. Combining this analysis with DIC microscopy, we were able to simultaneously assess the morphology of surface-wetted condensates and the formation of ThT-positive species.

As expected, increasing well surface hydrophobicity reduced the wettability of α-syn condensates. This was evident from the increased contrast at the condensate border and the more spherical morphology of surface-wetted condensates (Fig. 4a─c). Additionally, the increase in condensate diameter after wetting was significantly reduced on more hydrophobic surfaces (Supplementary Fig. 25), further confirming lower wettability.

**Figure 4.**
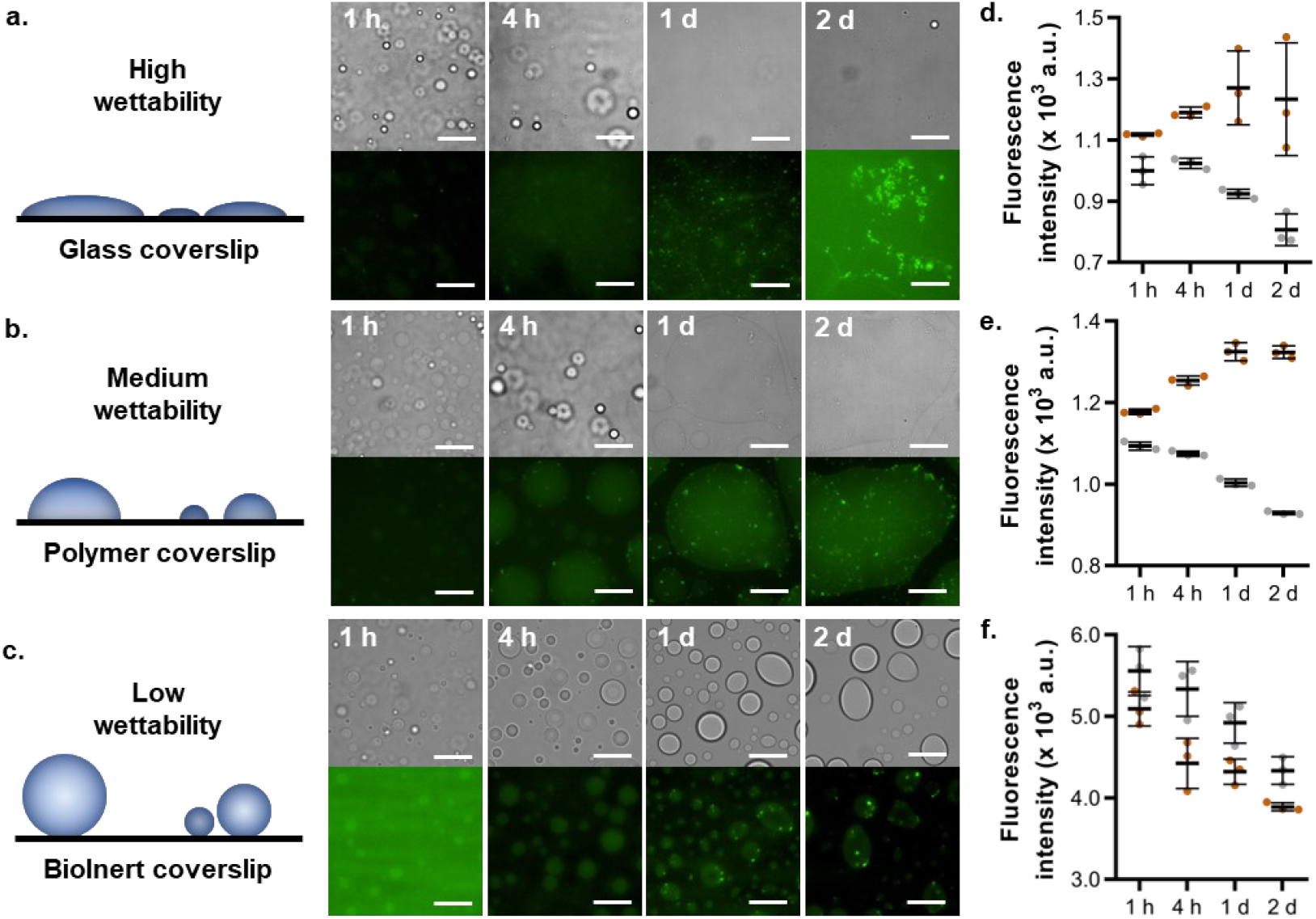
Wettability of FL α-syn on different surfaces under LLPS-promoting conditions. **(a─c)** Representative DIC (top) and fluorescent (bottom) images of 60 µM FL α-syn incubated with 25 µM PLK in LLPS buffer at 37 °C. Samples were incubated in 8-well slides with either a glass (a), polymer (b) or BioInert (c) coverslip. Images were acquired at selected time points at the bottom surface of the well. Scale bars represent 25 µm. **(d─f)** ThT fluorescence intensity quantification of the microscopy images acquired during the screening assay of glass (d), polymer (e) or BioInert (f) coverslips. Each data point is the average fluorescent intensity across the whole image, 3 images were quantified per time point. The mean intensity per time point is indicated by a solid black line, error bars represent standard deviation of the mean.

Over time, the number of small ThT-positive aggregates increased for all surface types. However, aggregates were observed earlier on the glass and polymer coverslips, with notably more formed by the end of this assay (Fig. 4a─c). This suggests that increased wettability accelerates aggregation, a trend further supported by the mean fluorescence intensity of the images, which showed a marked increase relative to the buffer control only for the glass and polymer surfaces (Fig. 4d─f and Supplementary Fig. 26). These findings highlight the importance of surface wetting in α-syn condensate maturation, providing evidence that the increased wettability of 11-140 and 19-140 α-syn likely underpins their accelerated LLPS-induced aggregation compared to the FL protein.

### The N-terminus regulates nucleation and propagation of α-syn fibrils

Our final aim was to analyze the microscopic steps of amyloid aggregation, so as to better understand the LLPS aggregation behavior of the truncated variants, particularly 5-140 α-syn. To monitor the kinetics of all three amyloid aggregation phases (*i*.*e*., the lag, growth, and plateau phases) in a single assay, in the absence of LLPS, it was necessary to incubate α-syn with ThT under aggregation promoting conditions, *i*.*e*., shaking in the presence of a glass bead at 37 °C (Fig. 5a and Supplementary Fig. 27). We then fitted the fluorescence data with a sigmoidal model to derive the time of the lag phase (t_lag_) and the growth rate (Hillslope) of aggregation. We found that all N-terminally truncated α-syn variants aggregated slower overall in a dispersed solution than the FL protein, in agreement with previous reports on similar protein variants.^34,36^ This behavior was mainly a consequence of significantly slower growth phases relative to FL α-syn, particularly 5-140 α-syn (Fig. 5a,b). In contrast, 5-140 α-syn had a significantly shorter lag phase than FL α-syn (~ 5.5-times faster t_lag_), while those of 11-140 and 19-140 α-syn were comparable (~ 1.5- and 1.1-times slower t_lag_, respectively) (Fig. 5a,c). Given the significant changes in the macroscopic phases of 5-140 α-syn aggregation, we decided to further characterize the aggregation mechanism. Thus, we monitored dispersed solution aggregation kinetics at varied FL and 5-140 α-syn concentrations, such that the scaling exponent (g), and thus dominant aggregation mechanism could be predicted using the AmyloFit software.^57^ We found that while FL α-syn displayed fragmentation dominated aggregation kinetics, *i*.*e*., a linear double logarithmic plot with g ~ 0.5, 5-140 α-syn exhibited an increase in the contribution of secondary nucleation, leading to fragmentation and secondary nucleation dominated aggregation kinetics (Supplementary Fig. 28).^57^

**Figure 5.**
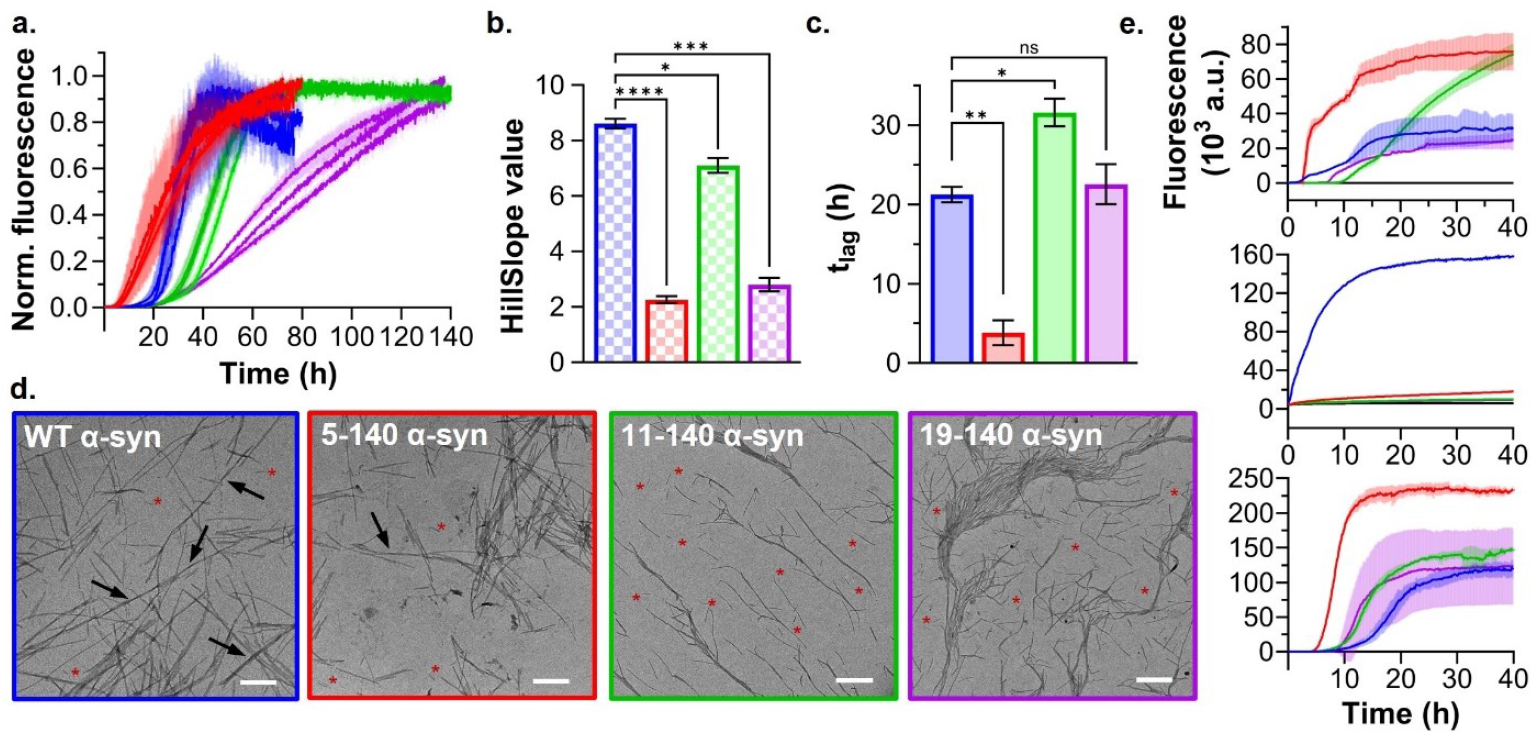
The microscopic steps of α-syn amyloid aggregation. **(a)** Dispersed solution aggregation of FL (blue), 5-140 (red), 11-140 (green) and 19-140 (purple) α-syn. Three individual biological repeats are shown. Each repeat is the mean of either two or three normalized technical replicates. The error bars represent the standard deviation of the mean. **(b/c)** Bar plots representing the Hillslope (b) and the t_lag_ (c) values of FL (blue), 5-140 (red), 11-140 (green) and 19-140 (purple) α-syn. Error bars represent the standard error of the mean of three biological repeats. **(d)** Representative TEM images of the insoluble fraction after aggregation. Short fibrils are highlighted by a red asterisk and fibril twists are indicated by an arrow. Scale bars represent 500 nm. **(e)** Aggregation kinetics of FL (blue), 5-140 (red), 11-140 (green) and 19-140 (purple) α-syn under conditions promoting lipid-induced primary nucleation (top), elongation (middle), or secondary nucleation (bottom). A single representative biological repeat is shown. Each repeat is the mean of three technical replicates. The semi-transparent error bars represent the standard deviation of the mean.

Subsequently, we quantified the percentage of monomeric protein converted into aggregates during the assay. We found similar amounts of soluble protein remained at the end of aggregation for all variants (~10–20 % of the initial monomer), indicating a similar conversion of monomers into aggregates (Supplementary Fig. 29). Finally, we characterized the morphology and size of the aggregates by performing TEM on the insoluble protein fractions isolated after the dispersed solution aggregation assay (Fig. 5d and Supplementary Fig. 30 and 31). Although all variants formed rod-like fibrils, 11-140 and 19-140 α-syn fibrils did not display any twisting.

Quantitative analysis of the TEM images highlighted that while FL and 5-140 α-syn fibrils were similar in length, 11-140 and 19-140 α-syn fibrils were typically shorter (Fig. 5d and Supplementary Fig. 30 and 31). Thus, increasing N-terminal truncation appears to reduce the structural stability of α-syn fibrils formed in a dispersed solution. This result is consistent with existing evidence that truncation of the first 6 N-terminal residues reduces fibril stability against mechanical agitation and protease activity.^34^ As the amyloids formed during our dispersed solution aggregation assay experience significant mechanical agitation due to shaking, our data indicate that deleting residues 5 and 6 destabilizes α-syn amyloids against mechanical stress-induced fragmentation.

5-140 and FL α-syn fibrils also shared a comparable width distribution, while both 11-140 and 19-140 α-syn amyloids were significantly thinner (Fig. 5d and Supplementary Fig. 30 and 31). Hence, 11-140 and 19-140 α-syn fibrils were likely at an earlier maturation stage, *i*.*e*., protofilaments and or protofibrils. This theory is supported by the high degree of fibrillar clumping observed for 11-140 and 19-140 α-syn, where many thin fibrils clustered together into large bundles (Fig. 5d and Supplementary Fig. 30). Such bundles may result from non-specific interactions between exposed hydrophobic regions on the protofilaments/protofibrils. Our findings are consistent with the inability of 7-140 α-syn to form thick fibrils, as we have previously reported,^34^ indicating deletion of residues 5 and 6 arrests dispersed solution aggregation at an earlier maturation stage.

Together, our data demonstrate that N-terminal truncation has a widespread effect on the dispersed solution amyloid aggregation of α-syn.

### The N-terminus regulates interaction of α-syn monomers with lipids and fibrils

To further our understanding of the role played by the N-terminus in the lag and growth phases of α-syn self-assembly, we examined the individual microscopic aggregation mechanisms, primary and secondary nucleation, and fibril elongation, for each of our variants.

First, we evaluated their propensity to undergo fibril elongation, *i*.*e*., monomer binding to the ends of pre-formed fibrils, by monitoring their aggregation at neutral pH in the presence of short pre-formed fibrils (1 min sonication, see Supplementary Methods).^58,59^ All N-terminally truncated variants demonstrated strongly inhibited fibril elongation with respect to FL α-syn (Fig. 5e). Thus, consistent with existing evidence,^36^ our data indicate that the first 4 residues of α-syn monomers are critical for their interaction with fibril ends, explaining the delayed growth phase observed during the dispersed solution aggregation assay.

To assess the ability of each variant to undergo heterogeneous primary nucleation, *i*.*e*., nucleation at the lipid membrane surface, we monitored their aggregation in the presence of 100 nm DMPS lipid vesicles.^60^ To instead investigate the secondary nucleation ability of the truncated variants, *i*.*e*., monomer nucleation at the surface of pre-formed fibrils, we monitored their aggregation at low pH in the presence of long pre-formed fibrils (15 s sonication, see Supplementary Methods).^58,61^ To ensure that all pre-formed fibrils were structurally mature and homogeneous, we used fibrils made of FL α-syn, as the aggregates of N-terminally truncated α-syn formed in a dispersed solution arrest at different maturation stages (Fig. 5d). The size homogeneity of all pre-formed fibril seeds and lipid vesicles was confirmed by DLS (Supplementary Fig. 32). Under both conditions, all α-syn variants aggregated, indicating N-terminally truncated monomers can interact with lipid vesicles and fibril surfaces to some extent (Fig. 5e).

11-140 and 19-140 α-syn displayed comparable aggregation kinetics, with inhibited lipid-induced primary nucleation and enhanced secondary nucleation, while both nucleation mechanisms were significantly accelerated for 5-140 α-syn (Fig. 5e). Given 5-140 α-syn lacks Asp2, decreased electrostatic repulsion towards the negatively charged DMPS lipid vesicles may promote primary nucleation. In contrast, the increased negative charge and absence of key lipid-binding residues in both 11-140 and 19-140 α-syn likely inhibit this nucleation^28^. Under secondary nucleation conditions, where the pH approaches α-syn’s isoelectric point and so electrostatic effects are minimized, the decreased hydrophobicity of all truncated variants may reduce hydrophilic-hydrophobic repulsion between α-syn monomers and the hydrophilic C-terminal fuzzy coat on fibrils, thus promoting secondary nucleation.

## Discussion

Here, we used a set of pathologically relevant N-terminal truncations to provide a detailed mechanistic description of α-syn LLPS and condensate maturation. To do so, we developed a strategy combining the use of microscopy and a microplate reader to monitor and quantify condensate formation, surface wetting, and amyloid aggregation. We found that while N-terminal truncation had little effect on α-syn condensate formation, the PTM significantly accelerated their maturation into amyloids (Fig. 3). In fact, a trend emerged: increasing N-terminal truncation, which reduces N-terminal hydrophobicity, delayed α-syn condensate sedimentation, increased condensate wettability on a hydrophilic well surface, and accelerated amyloid formation.

By modulating surface hydrophobicity, we demonstrated that increased wettability accelerated FL α-syn aggregation (Fig. 4). As one sample was prepared and then aliquoted across the different well slides, it is reasonable to assume that the total condensate volume is consistent across the slides. Consequently, increased wettability should result in a higher total condensate surface-to-volume ratio. Therefore, our data indicate that increasing the surface area of the condensate-bulk solution interface accelerates α-syn aggregation, implying the interface acts as a catalytic site for α-syn aggregation nucleation.

Given that coalescence between a condensate in solution and a surface-wetted condensate decreases the net surface-to-volume ratio, slower sedimentation will prolong a system of increased total surface-to-volume ratio. Combine this with their increased wettability and truncated α-syn condensates likely experience higher surface-to-volume ratios than those formed by the FL protein. Thus, the accelerated condensate maturation of N-terminally truncated α-syn could be explained by our interface-catalyzed nucleation model.

The notable exception to this trend was the pathological variant 5-140 α-syn. Despite having a more hydrophobic N-terminus than 19-140 α-syn, it exhibited an accelerated liquid-to-solid transition, with aggregate formation observed within condensates in solution (Fig. 3 and Supplementary Fig. 21). AmyloFit analysis of 5-140 α-syn dispersed solution aggregation kinetics predicted an increased contribution of secondary nucleation (Supplementary Fig. 28). This was confirmed by investigation into the microscopic aggregation mechanisms, which revealed significantly enhanced interface-catalyzed nucleation of 5-140 α-syn on both lipid vesicles and pre-formed fibrils (Fig. 5). Thus, 5-140 α-syn may aggregate faster within condensates because its nucleation at the condensate-bulk solution interface is also accelerated.

Interestingly, secondary nucleation was accelerated for all N-terminally truncated variants (Fig. 5). Given that PLK binds to and masks the negative C-terminus of α-syn, mimicking the effects of the low pH used to promote secondary nucleation, it is possible that enhanced secondary nucleation also contributes to the accelerated aggregation of truncated α-syn under LLPS conditions.

Overall, our data provides strong evidence that the interface of α-syn condensates catalyzes α-syn aggregation, consistent with existing evidence that α-syn can aggregate at the surface of pre-formed coacervates composed of inert biomolecules.^45^ In the context of neurodegeneration, changing α-syn condensate wettability within cells through therapeutic intervention could be a viable avenue towards mitigating α-syn toxicity. Our findings also indicate that α-syn LLPS and amyloid aggregation are linked, yet able to proceed independently. In fact, the same protein modification may have distinct effects on each pathway. This suggests that inhibitors of one process may not necessarily affect the other, a possibility that should be taken into consideration for future drug discovery. Our designed techniques establish a robust foundation for future investigations into the microscopic mechanisms mediating α-syn LLPS and amyloid aggregation.

## Materials and Methods

### Differential interference contrast and fluorescence microscopy

To assess the LLPS behavior of α-syn, the protein was incubated with PEG-8000 and PLK, conditions previously shown to induce α-syn LLPS.^42^ PEG-8000 was dissolved in PBS, pH 7.4 to a concentration of 20 % w/v. PLK hydrochloride with a molecular weight of 15,000–30,000 g/mol was dissolved in 10 mM HEPES, 100 mM NaCl, pH 7.4 to a concentration of 3 mM (determined using an approximate molecular weight of 22,500 g/mol). LLPS was induced by incubating monomeric protein (0, 20, 40 and 60 µM) with PLK (0, 25, 50 and 100 µM) in LLPS buffer (PBS, pH 7.4, 10 % PEG-8000, 0.02 % NaN_3_). 5 minutes after PLK addition, 100 µl of sample (single replicate per assay) was loaded into a 96 well full-area glass bottom plate (Sensoplate, Greiner Bio One, Austria). The plate was covered with a clear, microscopy compatible self-adhesive film (ibiSeal, Ibidi, Germany). The film was secured onto the plate using strips of adhesive aluminum foil along each edge. The samples were incubated at 37 °C within the chamber of a Nikon ECLIPSE Ti2-E microscope (Nikon, Japan) and automated DIC microscopy imaging in solution was carried out 15 min from PLK addition using a 40x air objective. A z-stack (3-step, 200 µm step size) within the bulk solution was acquired per well.

Condensate maturation in samples containing monomeric α-syn (0 or 60 µM) and PLK (0 or 25 µM) was monitored at 37 °C using the above automated solution imaging at pre-determined time points over a 20 h incubation period. Additionally, automated imaging was performed on the bottom surface of the well at each time point. Endpoint (bottom of well and in solution) DIC images were manually captured after the 20 h time point. When stated, 20 µM ThT was included to monitor amyloid fibril formation, and fluorescence images were taken alongside DIC images using excitation 405 nm and emission 515/30 nm filters. Unless otherwise stated a 40x air objective was used.

Both the 15 min and time course DIC solution images underwent processing, which involved estimating the number of objects per image and the area of every object in a given image. This analysis was conducted using GA3 software integrated into the NIS-Elements software (Nikon, Japan). A detailed GA3 recipe is provided in Fig. S4. In brief, images were first subjected to denoising to reduce shot noise. Objects were identified within the images by defining a gain thresholding range. To enhance accuracy, border objects and objects exceeding 50 µm^2^ in area (larger than the maximum condensate area determined through manual inspection) were excluded. The compiled object area values and the total number of objects per image, across all z positions, per well, and time point were consolidated into a single data table.

Phase diagrams were generated using the normalized total number of objects counted per well, summed over all z positions, 15 min from PLK addition. To examine the time course experiments, the area distribution of each sample, compiled over all z positions, at each time point was plotted. The mean area at each time point was determined, plotted against time, and normalized. Three separate biological repeats were globally fitted with a one-phase association curve. The half-time (t_50_) of object growth was derived. Next, the total object count at each time point, compiled over all z positions, was plotted against time and normalized. Three separate biological repeats were globally fitted with a one-phase decay curve and the half-time (t_50_) of count decay was derived. Normalization of the object count and mean object area values was performed to account for inherent variations in their absolute values arising from manual error when defining the thresholding range. All data were plotted using GraphPad Prism version 10.0.3.

FL α-syn localization was monitored using a 1:100 ratio of rhodamine labeled: unlabeled FL α-syn. Fluorescence images were taken alongside DIC images using excitation 525 nm and emission 641/75 nm filters. A single sample was prepared at a time, loaded into a 96 well full-area glass bottom plate, and imaged manually using a 40x air objective.

### Aggregation assay under liquid-liquid phase separation conditions

Fibril formation was monitored by incubation of monomeric α-syn (0 and 60 µM) with PLK (0 and 25 µM) and 20 µM ThT in LLPS buffer. 100 µl of sample (3 replicates) was loaded into a 96 well full-area glass bottom plate, sealed with aluminum foil, and incubated at 37 °C under quiescent conditions for ~ 70 h in a CLARIOstar Plus microplate reader. Fluorescent intensity measurements were taken at the center of the well, excitation 440-10 nm, dichroic 460 nm and emission 480-10 nm filters, 2 gains and 30 flashes per well, every 20 min. All sample spectra were background corrected by subtracting the spectrum of 20 µM ThT in LLPS buffer, with data extracted from the same assay background corrected using the same buffer spectrum. The data were plotted using GraphPad Prism version 10.0.3.

### Liquid-liquid phase separation surface hydrophobicity assay

Surface wetting and aggregate formation was monitored by incubation of 0 or 60 µM FL α-syn with 25 µM PLK and 20 µM ThT in LLPS buffer. 200 µl of sample was loaded into µ-slide 8-well^high^ (Ibidi, Germany) with either a glass coverslip, a polymer coverslip, or a BioInert polymer coverslip. Samples were incubated at 37 °C for 2 d and the bottom of the well imaged at selected time points by DIC and ThT fluorescence microscopy (excitation 405 nm, emission 515/30 nm filters) using a 60x oil objective. The mean fluorescence intensity of each image was measured and plotted using GraphPad Prism version 10.0.3.

## Acknowledgments

We thank the UK Research and Innovation (Future Leaders Fellowship MR/S033947/1 and Fellowship renewal MR/Y003616/1), the Engineering and Physical Sciences Research Council (grant EP/S023518/1), Alzheimer’s Society, UK (grant 511), Alzheimer’s Research UK (ARUK-PG2019B-020) and the Department of Chemistry of Imperial College London (PhD scholarship).

We also thank Prof. Dr. Paolo Arosio (ETH Zurich) and Prof. Annalisa Pastore (Imperial College London) for helpful discussions and feedback on the manuscript. Furthermore, we thank the Chemistry Mass Spectrometry facilities for assistance with ESI-MS, and the Electron Microscopy Centre facilities at The Center of Structural Biology for assistance with TEM. Fig. 1a includes a darkened version of the multiwell-plate-3d icon by Servier.

## Author contributions

RJT, DMV and SCA performed all experiments. RJT and FAA analyzed the data. RJT and FAA conceptualized the work. All authors discussed the data and provided feedback on the manuscript.

