## Supplementary Information for "Surface Wetting Is a Key Determinant of α-Synuclein Condensate Maturation"

**This PDF file includes:**

Supplementary Methods
Supplementary Table 1 and 2
Supplementary Figures 1 to 32
Supplementary References

**Other Supplementary Information for this manuscript includes:**

Supplementary Movies 1 to 4

### Extended Materials and Methods

No unexpected or unusually high safety hazards were encountered.

#### Protein expression and purification

FL α-syn (residues 1-140) was expressed and purified as detailed previously.[1] Briefly, pT7-7 α-syn FL plasmid (a gift from Hilal Lashuel, Addgene, USA[2]) was transformed into BL21-Gold (DE3) competent Escherichia coli (E. coli) cells (Agilent Technologies) according to the manufacturer's instructions and previous methodology. Transformed cells were grown (37 °C, 200 rpm shaking) in LB media containing ampicillin (100 µg/ml) to an OD600 of ~ 0.7 before FL α-syn expression induced with 1 mM IPTG overnight (28 °C, 200 rpm shaking). The cells were then harvested by centrifugation and resuspended in buffer A (20 mM Tris-HCl, 1 mM EDTA, pH 8.0) including a protease inhibitor tablet (EDTA-free). Cells were lysed by sonication on ice before cell debris was removed by centrifuged (18,000 rpm, 45 min). The lysate was then boiled at 80 °C for 20 min, followed by another round of centrifugation (18,000 rpm, 30 min). Next, streptomycin sulphate (10 mg/ml) was gradually introduced to the supernatant. The mixture was incubated with rotation before further centrifugation (18,000 rpm, 30 min). FL α-syn was precipitated by the slow addition of ammonium sulphate (360 mg/ml) on ice, followed by incubation with rotation, and a final centrifugation (18,000 rpm, 30 min). The pellet was collected, re-suspended and dialyzed into buffer A overnight. The sample was then loaded onto an anion exchange chromatography column (HiPrep Q HP 16/10, Cytiva, USA) and further purified using a gradient elution with buffer B (20 mM Tris-HCl, 1 M NaCl, 1 mM EDTA, pH 8.0). The FL α-syn protein fractions, as determined by sodium dodecyl sulphate–polyacrylamide gel electrophoresis (SDS-PAGE), were then dialyzed into phosphate buffered saline (PBS, pH 7.4) overnight, before being passed through a gel filtration column (HiLoad 26/600 Superdex 75 pg, Cytiva, USA). The pure fractions, determined by SDS-PAGE, were combined and the concentration determined by Ultraviolet–visible (UV-Vis) spectroscopy absorbance at 275 nm ($\varepsilon_{275nm}$ = 5600 $M^{-1}cm^{-1}$).

To generate 11-140 and 19-140 α-syn, deletion polymerase chain reaction (PCR) was performed on the pT7-7 Int7-140αSyn plasmid, which encodes a protein sequence where an intein is fused to the N-terminus of residues 7-140 of α-syn,[1] eliminating the need for a non-native starting Met in the protein sequences. PCR was carried out using the Q5 high-fidelity DNA polymerase, the 5' phosphorylated primer pairs and their corresponding annealing temperatures ($T_a$), listed in Table S1. The template DNA was then digested with DpnI. Following sample purification using a NucleoSpin Gel and PCR Clean-up kit (MACHEREY-NAGEL, Germany), T4 DNA ligase was used to ligate the 5' phosphorylated blunt-end DNA fragments. All steps were carried out according to the relevant manufacturer's instructions. The resultant plasmids were then transformed into XL10-Gold Ultracompetent E. coli cells and purified using a QIAprep Spin Miniprep Kit. Mutagenesis success was determined by sequencing (GENEWIZ). Following several failed attempts to generate the 11-140 α-syn plasmid, Genscript performed the mutagenesis on our behalf.

Both variant plasmids were then transformed into BL21-Gold (DE3) cells, we found that increasing initial incubation on ice from 30 to 90 min increased transformation efficiency. The intein-α-syn fusion proteins corresponding to 11-140 and 19-140 α-syn were then expressed and the cell lysate harvested as described for FL α-syn. Subsequently, the fusion proteins were immobilized on a chitin resin before the target α-syn variant was cleaved, eluted and further purified by size-exclusion chromatography as previously described.[1]

The N-terminal residue of 5-140 α-syn is Met and so the variant does not require intein conjugated expression. To generate the mutated plasmid the above deletion PCR process was applied to the pT7-7 α-syn WT plasmid using the 5' phosphorylated primer pair listed in Table S1. 5-140 α-syn was then expressed and purified as detailed for the FL protein.

#### Fluorescent labelling

FL α-syn was covalently tagged at its lysine residues using NHS-Rhodamine (ex. 552 nm, em. 575 nm). The probe was added to 100 µM α-syn in ~ 8-fold molar excess, and the reaction was allowed to proceed at RT (~ 4 h, gentle mixing). Subsequently, excess dye was removed by extensive dialysis into PBS, pH 7.4 (overnight, 4 °C). For the purpose of subsequent assays, it was sufficient to assume no significant change in α-syn concentration had occurred during this process.

Electrospray ionization mass spectrometry

Purified protein samples (~ 50 µM, diH2O) were analyzed by electrospray ionization mass spectrometry (ESI-MS). ESI-MS was performed by Lisa Haigh, Malgorzata Puchnarewicz and Adriana Lobosco using the Chemistry Mass Spectrometry facilities available at the Molecular Sciences Research Hub, Department of Chemistry, Imperial College London. The spectra were plotted using GraphPad Prism version 10.0.3 (GraphPad Software).

#### Transmission election microscopy

A sample of the washed insoluble fraction taken at the endpoint of a dispersed solution aggregation assay, or of the whole solution taken at the endpoint of an aggregation assay under LLPS conditions, was applied to carbon films on 300 mesh copper grids (Agar Scientific Ltd., UK). The grids were quickly washed with diH2O, negatively stained with 2 % (w/v) uranyl acetate and washed again. The grids were imaged using the Tecnai 12 Spirit transmission electron microscope (Thermo Fisher Scientific (formerly FEI), USA) available at the Electron Microscopy Centre, Centre of Structural Biology, Imperial College London. Where specified fibril lengths and widths were measured using Fiji, with the data plotted using GraphPad Prism version 10.0.3.

#### Dot-blot analysis

The presence of α-syn in the insoluble aggregates formed following aggregation under LLPS conditions was determined by dot-blot analysis. The soluble and insoluble protein fractions at the endpoint of the aggregation assay were separated by centrifugation (30 min, 14,000 rpm). The insoluble pellet was resuspended in a minimum volume of PBS before 5 replicates of 3 µl aliquots were transferred to a nitrocellulose membrane (0.45 µm) and allowed to dry. The membrane was then blocked (1 h, 4 °C) with a solution of milk powder (5 %) dissolved in PBS-T (PBS supplemented with 0.1 % Tween 20). After washing with PBS-T, the membrane was incubated (overnight, 4 °C) with the anti-α-syn primary antibody MJFR1 (diluted 1:1000 in PBS-Tween, Abcam, UK). Following rigorous washing with PBS-T, the membrane was incubated (1 h, RT) with the goat anti-rabbit IgG (H + L) highly cross adsorbed secondary antibody, conjugated to the fluorophore Alexa Fluor plus 555 (diluted 1:5000 in PBS-Tween, Thermo Fisher Scientific, USA). Finally, the membrane was washed with PBS-T and imaged with a Typhoon FLA 9000 (Cytiva, USA). As a control, the same dot-blot protocol was used to analyze the reactivity of the primary antibody against 25 µM PLK in PBS.

#### Circular dichroism

The far-UV circular dichroism (CD) spectra of fresh monomeric protein (20 µM) or the insoluble aggregate pellet (~ 10 µM) taken from the endpoint of a ThT aggregation assay under LLPS conditions were recorded. The insoluble fractions were obtained by centrifugation, and their concentration estimated using UV-Vis spectroscopy absorbance at 275 nm ($\varepsilon_{275nm}$ = 5600 $M^{-1}cm^{-1}$) following denaturation in 4 M guanidinium chloride for 2 h at RT. Spectra of all samples were taken using a Chirascan v100 (Applied Photophysics Ltd, UK) from 200 to 250 nm with a 0.5 nm step, 1 nm bandwidth, 1 s per time point and 5 accumulations. A PBS background spectrum was subtracted from each sample spectrum, with data extracted from the same assay background corrected using the same buffer spectrum. For monomeric protein samples the raw data (in mdeg) were converted to mean residue ellipticity (MRE, units deg $cm^2$ $dmol^{-1}$) using:[3]

$$MRE = mdeg \div (l \times c \times (n-1)) \quad \text{equation (1)}$$

where mdeg is the raw data (in mdeg), l is the cuvette pathlength (in mm), c is the sample concentration (in M) and n is the number of amino acids. The data were plotted using GraphPad Prism version 10.0.3.

#### Dispersed solution aggregation assay

To monitor all three phases in the amyloid aggregation, monomeric protein was aggregated and monitored by fluorescence spectroscopy using a previously described protocol.[1] Briefly, monomer solutions (50 µM) were aggregated with ThT (20 µM) and $NaN_3$ (0.02 %) in PBS (pH 7.4). 170 µl of sample (in triplicate) was loaded into a 96 well full-area µClear plate (non-binding, clear bottomed, #655906, Greiner Bio-One, Austria), sealed with an aluminum film and incubated at 37 °C for ~ 70 - 140 h in a CLARIOstar Plus microplate reader (BMG Labtech, Germany). Aggregation was promoted by linear shaking (300 rpm, 300 s before each cycle) and addition of a single borosilicate bead (3 mm diameter) per well. Fluorescent intensity measurements were taken every 520 s using spiral averaging (5 mm diameter) and excitation 440 nm, dichroic 460 nm, and emission 480 nm filters, 3 gains and 50 flashes per well. All sample spectra were background corrected by subtracting the spectrum of buffer alone (i.e., 20 µM ThT and 0.02 % $NaN_3$ in PBS), with data extracted from the same assay background corrected using the same buffer spectrum.

The data were plotted using GraphPad Prism version 10.0.3. The individual biological repeats were fit with sigmoidal, 4PL, X is concentration standard curves provided by the software. $t_{50}$ was estimated as the time (x-value) at which 50 % maximum fluorescence intensity was reached. $t_{lag}$ was estimated by extrapolation of the tangent at $t_{50}$. The steepness of the curve was estimated by the Hillslope value, where Prism assigns a standard sigmoidal receptor binding curve a Hillslope value of 1.0, while a steeper curve has a higher Hillslope value, and a shallower curve has a lower Hillslope value.

To estimate ɣ for FL and 5-140 α-syn, this assay was repeated with 10, 25, 50, 75 and 100 µM protein. The data were uploaded to AmyloFit 2.0 where the log of initial monomer concentration (m0) was calculated alongside the log of the half-life of aggregation ($t_{50}$).[4] All data were plotted using GraphPad Prism. The double logarithmic plot was fitted with a simple linear regression and ɣ estimated as the gradient. As ɣ is not constant for 5-140 α-syn, the data was fit with two straight lines corresponding to two different ɣ values.

### Lipid-induced aggregation assay

Lipid vesicles were produced based on a previously described protocol.[5] Briefly, 1,2-dimyristoyl-sn-glycero-3-phospho-L-serine (sodium salt) was dissolved in 20 mM phosphate buffer (PB), pH 6.5 at ~ 70 °C for 2 h. The lipids were then freeze-thawed using liquid nitrogen and a ~ 50 °C water bath 5 times before being extruded 11 times through a membrane (100 nm pore diameter) at ~ 70 °C. Subsequently, monomer solutions (50 µM) were aggregated in PB (pH 6.5) in the presence of DMPS lipid vesicles (250 µM), ThT (20 µM) and $NaN_3$ (0.02 %). 170 µl of each sample (3 replicates) was loaded into a 96 well full-area µClear plate, sealed with aluminum foil, and incubated at 37 °C for ~ 40 h in a FLUOstar Omega microplate reader (BMG Labtech, Germany). Fluorescent intensity measurements were taken using spiral averaging (3 mm diameter), excitation 440-10 nm, dichroic 460 nm and emission 480-10 nm filters, 4 gains and 50 flashes per well. All sample spectra were background corrected by subtracting the spectrum of buffer alone (i.e., 20 µM ThT and 0.02 % $NaN_3$ in PB), with data extracted from the same assay background corrected using the same buffer spectrum. The size homogeneity of the lipid vesicles was determined by dynamic light scattering (DLS). All data were plotted using GraphPad Prism version 10.0.3.

### Seeded aggregation assay

Pre-formed fibril seeds were generated by shaking (~ 600 rpm) monomeric FL α-syn (~ 200 µM, PBS, pH 7.4) at 37 °C for ~ 5–10 days. $NaN_3$ (0.02 %) was added to prevent bacterial growth during prolonged incubation. The soluble and insoluble protein fractions were then separated by centrifugation (30 min, 14,000 rpm), the soluble fraction removed, and the pellet resuspended in fresh PBS, pH 7.4. Centrifugation and pellet resuspension were repeated before the final protein concentration was estimated using UV-Vis spectroscopy absorbance at 275 nm ($\varepsilon_{275nm}$ = 5600 $M^{-1}cm^{-1}$) after denaturation in 4 M guanidinium chloride for 2 h at RT.

Seeded aggregation assays were carried out based on previously described protocols.[6,7] Immediately before use the pre-formed FL α-syn fibril stock was diluted to ~ 100 µm in PBS (pH 7.4) and the precise concentration measured by UV-Vis spectroscopy. When performing a fibril elongation assay, this pre-formed fibril solution was then diluted to 50 µM in PBS, pH 7.4 and probe sonicated on ice at 20 % power, 15 s pulse, 15 s rest for 4 cycles. Monomeric protein (50 µM) was then incubated with these short pre-formed FL α-syn fibrils (5 µM), ThT (20 µM) and $NaN_3$ (0.02 %) in PBS, pH 7.4.

To perform secondary nucleation experiments, the ~ 100 µm pre-formed fibril stock was diluted to 5 µM in 20 mM acetic acid (NaCl 150 mM, pH 4.8) and probe sonicated on ice at 20 % power, 5 s pulse, 5 s rest, 3 cycles. Monomeric protein (50 µM) was also buffer exchanged into 20 mM acetic acid (NaCl 150 mM, pH 4.8), and then incubated with these long pre-formed FL α-syn fibrils (50 nM), ThT (20 µM) and $NaN_3$ (0.02 %).

For both assays, 170 µl of sample (3 replicates) was loaded into a 96 well full-area µClear plate, sealed with aluminum foil and incubated for ~ 40 h at 37 °C in a FLUOstar Omega microplate reader under quiescent conditions. Fluorescent intensity measurements were taken using the settings described above for the lipid-induced aggregation assay. All fibril elongation or secondary nucleation sample spectra were background corrected by subtracting the spectrum of buffer alone (i.e., 20 µM ThT and 0.02 % $NaN_3$ in either PBS or 20 mM acetic acid, respectively), with data extracted from the same assay background corrected using the same buffer spectrum. The size homogeneity of any sonicated pre-formed fibril seeds was determined by DLS. All data were plotted using GraphPad Prism version 10.0.3.

### Monomer conversion analysis

The soluble and insoluble protein fractions before and after a dispersed solution aggregation assay were separated by centrifugation (30 min, 16,900 x g). The soluble fraction was then extracted and analyzed by SDS-PAGE. For each sample, the mean grey value of the band corresponding to the relevant monomeric protein variant was measured using Fiji.[8] Percentage conversion of monomer into insoluble aggregates was estimated by comparison of the band intensity before and after aggregation. The data were plotted using GraphPad Prism version 10.0.3.

### Statistical analysis

Statistical significance was performed on the $t_{lag}$ and Hillslope values extracted from the dispersed solution aggregation assay data. A Welch and Brown-Forsythe ANOVA with multiple comparisons against the control (FL α-syn) was used where $^{ns}p \geq 0.05$, $0.01 \leq {}^{*}p < 0.05$, $0.001 \leq {}^{**}p < 0.01$, $0.0001 \leq {}^{***}p < 0.001$ and $^{****}p < 0.0001$.

Statistical significance was also performed on the half-life ($t_{50}$) of the decay in object count values extracted from the DIC time course experiments under LLPS conditions. A Welch and Brown-Forsythe ANOVA with multiple comparisons against the control (FL α-syn) was used with the same p values listed above.

**Table S1.**

5’ phosphorylated primer pairs, shown in the 5’ to 3’ direction, used to mutate the FL α-syn gene.

| α-Syn variant | Primer Pair | $T_a$ / °C |
|---|---|---|
| 5-140 | Forward: aaaggactttcaaaggcc<br>Reverse: catatgtatatctccttcttaaagttaaac | 60 |
| 11-140 | Forward: gccaaggagggagttgtg<br>Reverse: gttctgtacaacaacctgagatcc | 66 |
| 19-140 | Forward: gctgagaaaaccaaacagggtgtg<br>Reverse: gttctgtacaacaacctgagatccaagc | 70 |

**Table S2.**

Schematic representation of the N-terminally truncated α-syn variants selected for analysis[1].

| α-Syn variant | ΔCharge | Hydrophobicity* | N-terminus |
|---|---|---|---|
| WT (1-140) | 0 | 6.4 | $H_3N$-MDVFMKGLSKAKEGVVAAAE |
| 5-140 | +1 | 1 | $H_3N$-MKGLSKAKEGVVAAAE |
| 11-140 | -1 | 4.3 | $H_3N$-AKEGVVAAAE |
| 19-140 | -1 | -1.7 | $H_3N$-AE |

1. The first 20 residues of FL α-syn are shown with neutral residues in black, positive residues in green, negative residues in red. Subsequently, N-terminally truncated variants 5-140, 11-140 and 19-140 α-syn are shown with their corresponding residues deleted. Resultant change in charge (ΔCharge) and hydrophobicity for each truncated variant, relative to the FL protein, are shown. *Calculated according to Kyte-Doolittle scale.

**a.**

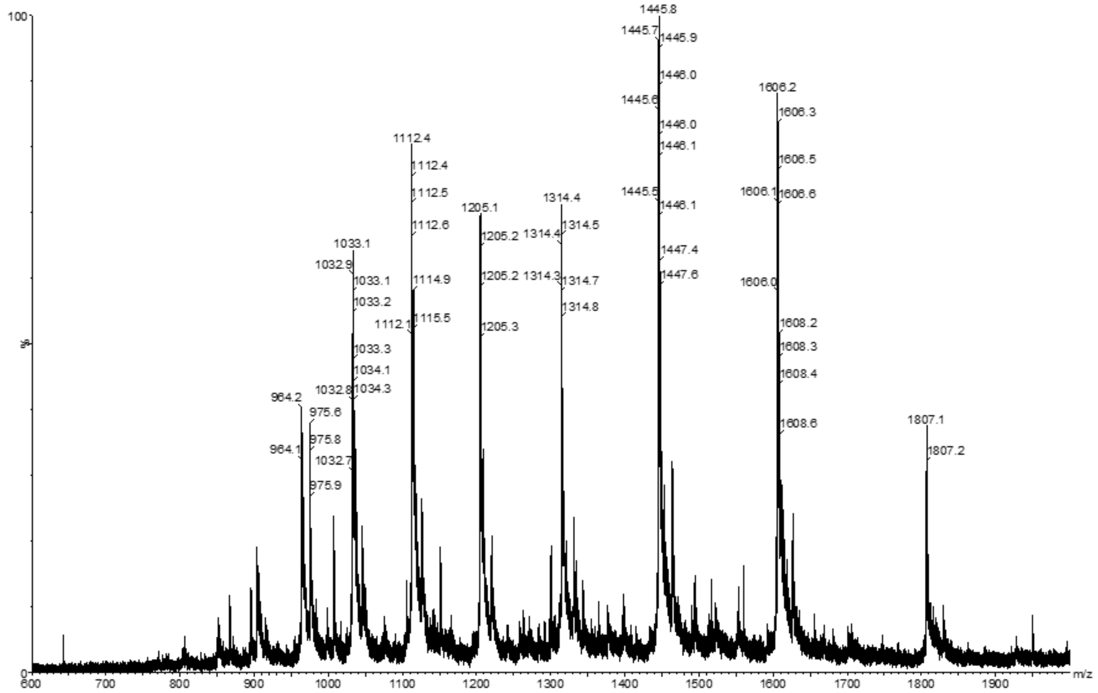
100
%
0
1445.8
1445.7
1445.9
1446.0
1445.6
1446.0
1446.1
1445.5
1446.1
1447.4
1447.6
1606.2
1606.3
1606.5
1606.1
1606.6
1606.0
1608.2
1608.3
1608.4
1608.6
1112.4
1112.4
1112.5
1112.6
1114.9
1112.1
1115.5
1205.1
1205.2
1205.2
1205.3
1314.4
1314.4
1314.5
1314.3
1314.7
1314.8
1033.1
1032.9
1033.1
1033.2
1033.3
1034.1
1032.8
1034.3
964.2
975.6
975.8
964.1
1032.7
975.9
1807.1
1807.2
600
700
800
900
1000
1100
1200
1300
1400
1500
1600
1700
1800
1900
m/z

**b.**

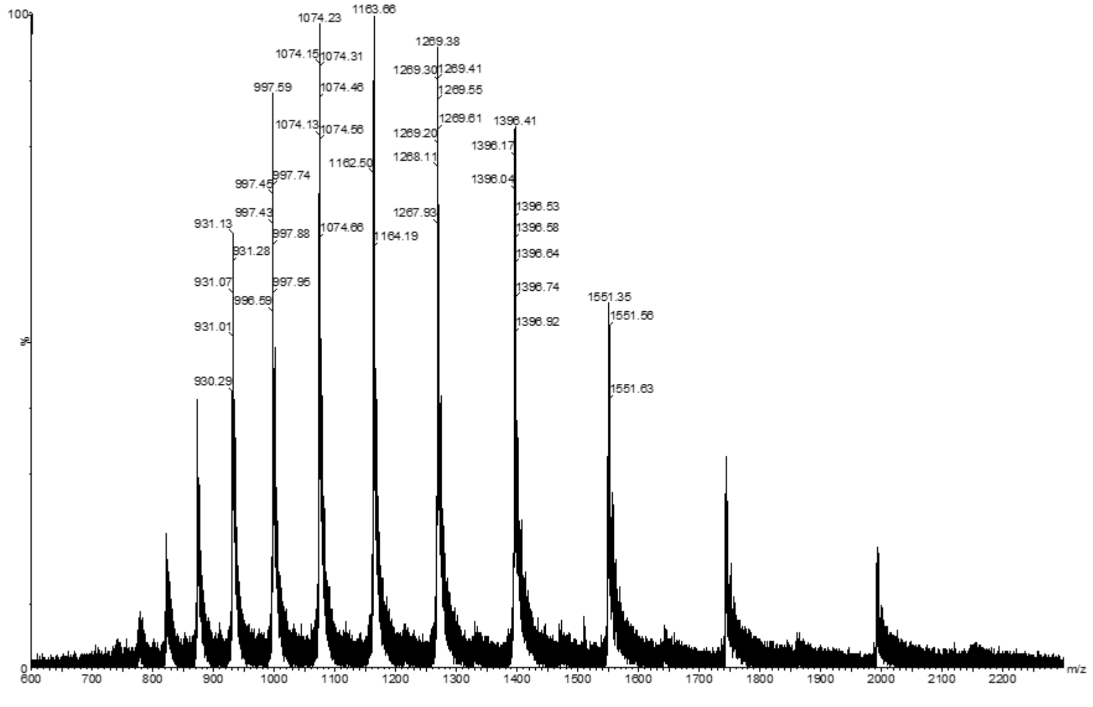
100
%
0
1074.23
1163.66
1074.15
1074.31
997.59
1074.46
1074.13
1074.56
997.74
997.45
997.43
931.13
997.88
1074.66
931.28
931.07
997.95
996.59
931.01
930.29
1162.50
1164.19
1269.38
1269.30
1269.41
1269.55
1269.61
1269.20
1268.11
1267.93
1396.41
1396.17
1396.04
1396.53
1396.58
1396.64
1396.74
1396.92
1551.35
1551.56
1551.63
600
700
800
900
1000
1100
1200
1300
1400
1500
1600
1700
1800
1900
2000
2100
2200
m/z

c.

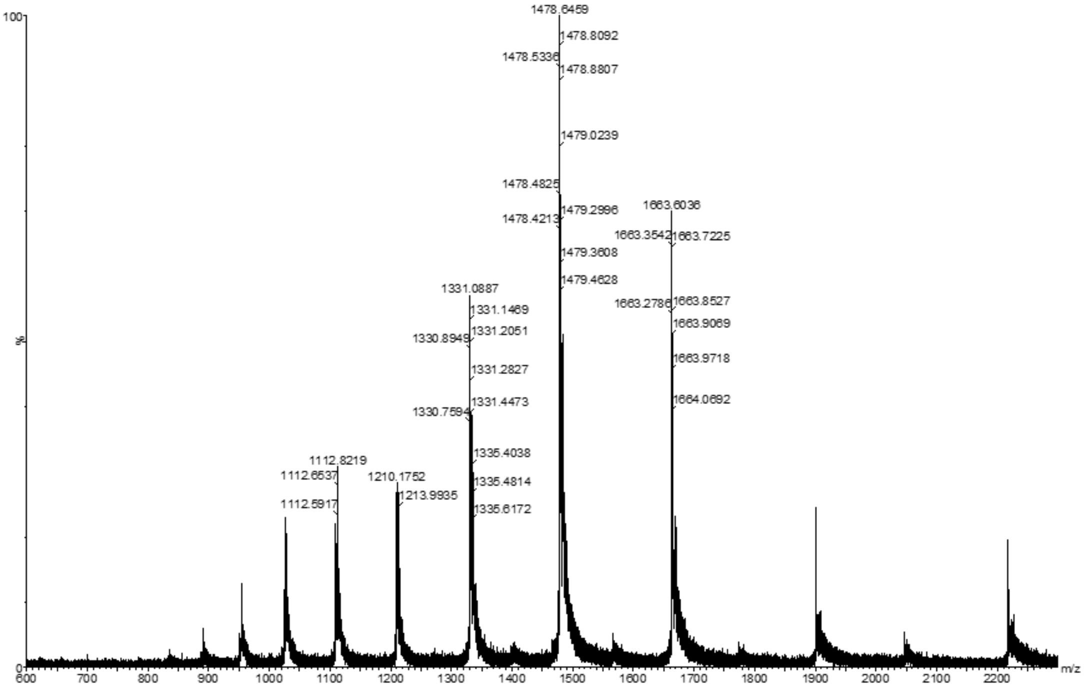
100
%
0
1478.6459
1478.8092
1478.5336
1478.8807
1479.0239
1478.4825
1479.2996
1478.4213
1663.6036
1663.3542
1663.7225
1479.3608
1479.4628
1331.0887
1331.1469
1663.2786
1663.8527
1663.9069
1330.8949
1331.2051
1663.9718
1331.2827
1331.4473
1664.0692
1330.7594
1112.8219
1335.4038
1112.6537
1210.1752
1335.4814
1112.5917
1213.9935
1335.6172
600 700 800 900 1000 1100 1200 1300 1400 1500 1600 1700 1800 1900 2000 2100 2200
m/z

d.

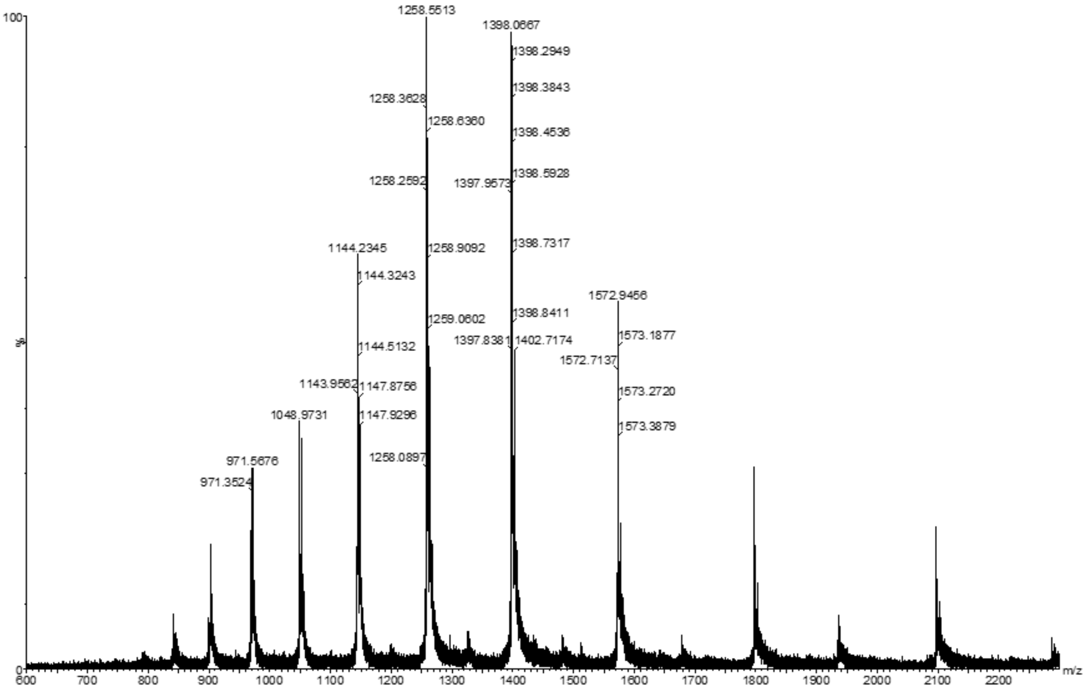
100
%
0
1258.5513
1398.0667
1398.2949
1398.3843
1258.3628
1258.6360
1398.4536
1258.2592
1397.9573
1398.5928
1144.2345
1258.9092
1398.7317
1144.3243
1572.9456
1259.0602
1398.8411
1144.5132
1397.8381
1402.7174
1573.1877
1572.7137
1143.9562
1147.8756
1573.2720
1048.9731
1147.9296
1573.3879
1258.0897
971.5676
971.3524
600 700 800 900 1000 1100 1200 1300 1400 1500 1600 1700 1800 1900 2000 2100 2200
m/z

e.

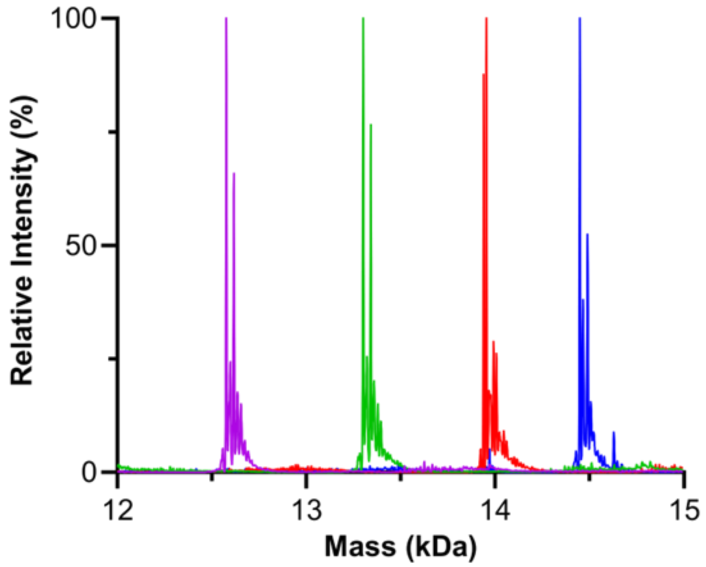


**Figure S1. N-terminally truncated α-syn purification confirmed by ESI-MS.**

a-d, ESI-MS spectra of FL (a), 5-140 (b), 11-140 (c) and 19-140 (d) α-syn prior to deconvolution. e, Deconvoluted ESI-MS spectra of FL (blue), 5-140 (red), 11-140 (green) and 19-140 (purple) α-syn. Experimental and expected masses, respectively, are as follows; FL α-syn 14449.0 Da and 14451.2 Da, 5-140 α-syn 13954.0 Da and 13959.0 Da, 11-140 α-syn 13302.0 Da and 13314.6 Da, 19-140 α-syn 12577.0 Da and 12589.2 Da.

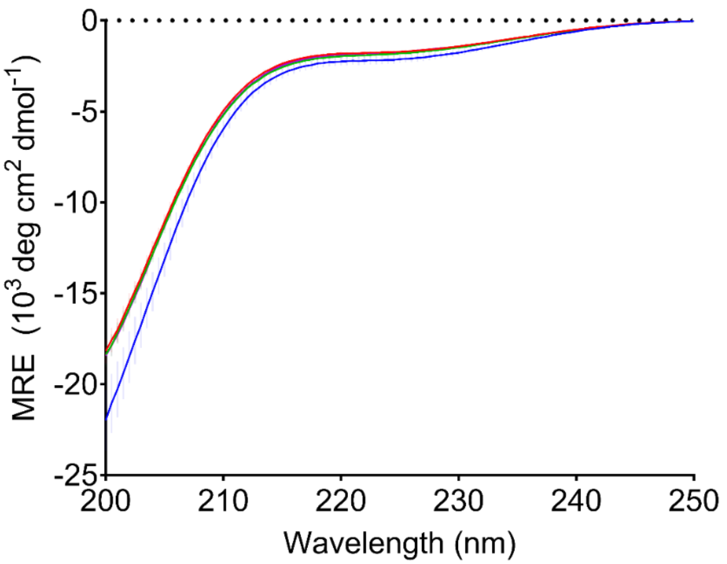


**Figure S2. N-terminally truncated α-syn retains random coil secondary structure.**

Far-UV CD spectra of monomeric FL (blue), 5-140 (red), 11-140 (green) and 19-140 (purple) α-syn. The mean of three individual biological repeats is shown per sample, semi-transparent error bars represent the standard error of the mean.

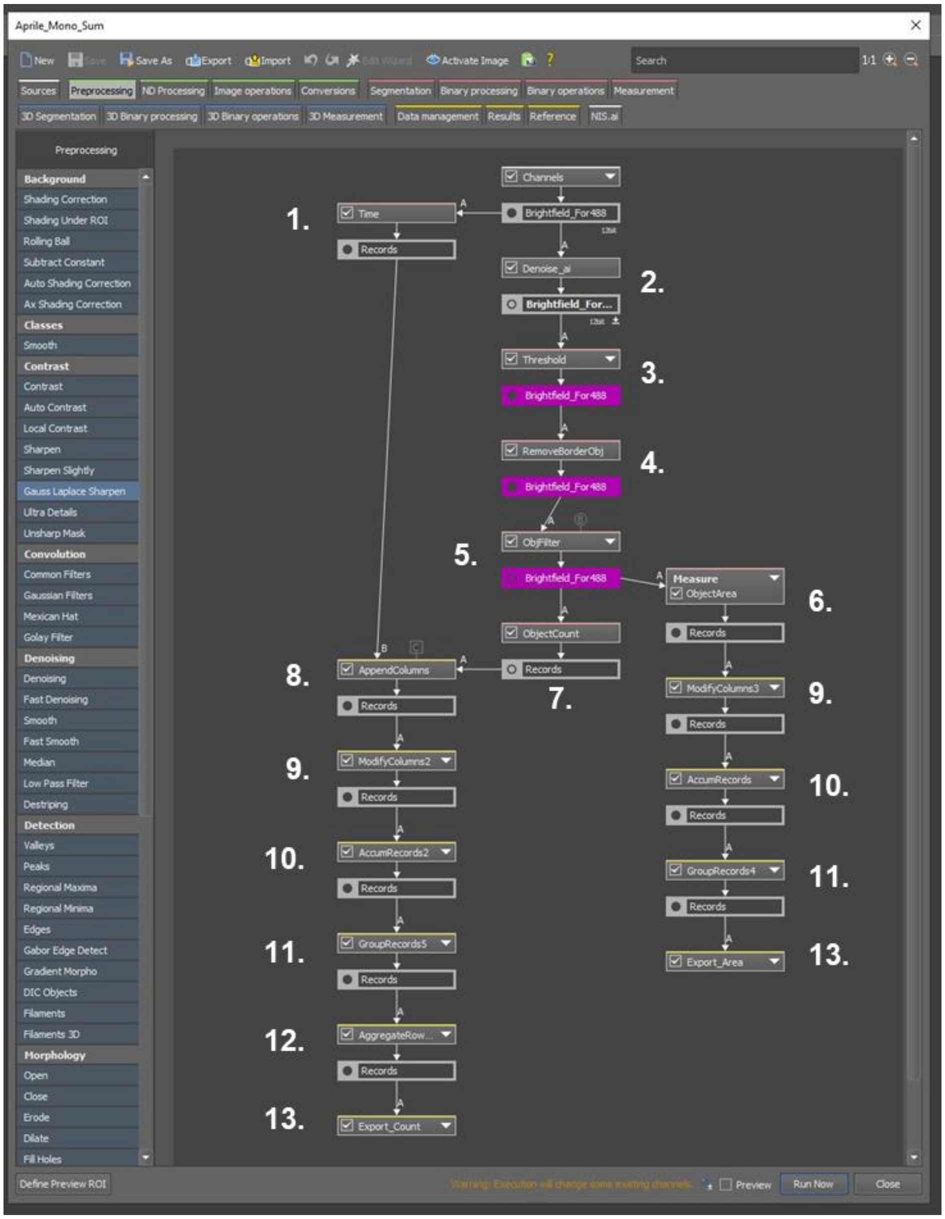


**Figure S3. GA3 recipe processes DIC images, quantifying object count and area.**

Step 1 acquires the time (in s) each image was taken. Step 2 runs denoise software on the DIC images. Step 3 requires manual setting of the gain threshold range used for object identification. Step 4 reduces object misidentification by removing any objects that are touching the image boarders. Step 5 further reduces object misidentification by removing any objects that are over 50 µm2 in area. Step 6 measures the area of every object within an image. Step 7 counts the number of objects per image. Step 8 compiles the object count and time stamp data into a single record. Step 9 removes any unwanted data from records. Step 10 repeats the object number count and object area measurements in every image taken (over every well position, z position and timepoint) and compiles them into the records. Step 11 orders and groups the records based on well position and then timepoint. Step 12 sums the total number of objects over the three z-stack images taken per well position and time point. Step 13 exports the record as a CSV file. It must be noted that this analysis was aided by visual inspection of the DIC images as this recipe cannot distinguish between condensates and other objects, e.g., dust particles. However, given the large number of condensates relative to aggregates/impurities, our quantitative analysis is robust.

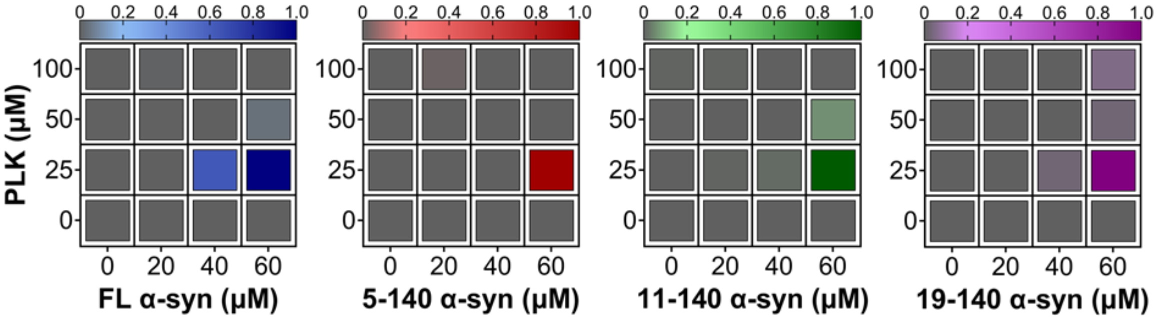


**Figure S4. N-terminal truncation does not affect α-syn condensate formation.**

Representative phase diagrams showing the normalized total object count at different protein and PLK concentrations, as determined by DIC images taken at 37 °C, 15 mins after PLK addition to FL (blue), 5-140 (red), 11-140 (green), and 19-140 (purple) α-syn in LLPS buffer. Object count values for each time point are the normalized sum of the individual object counts of the three z-stack images acquired.

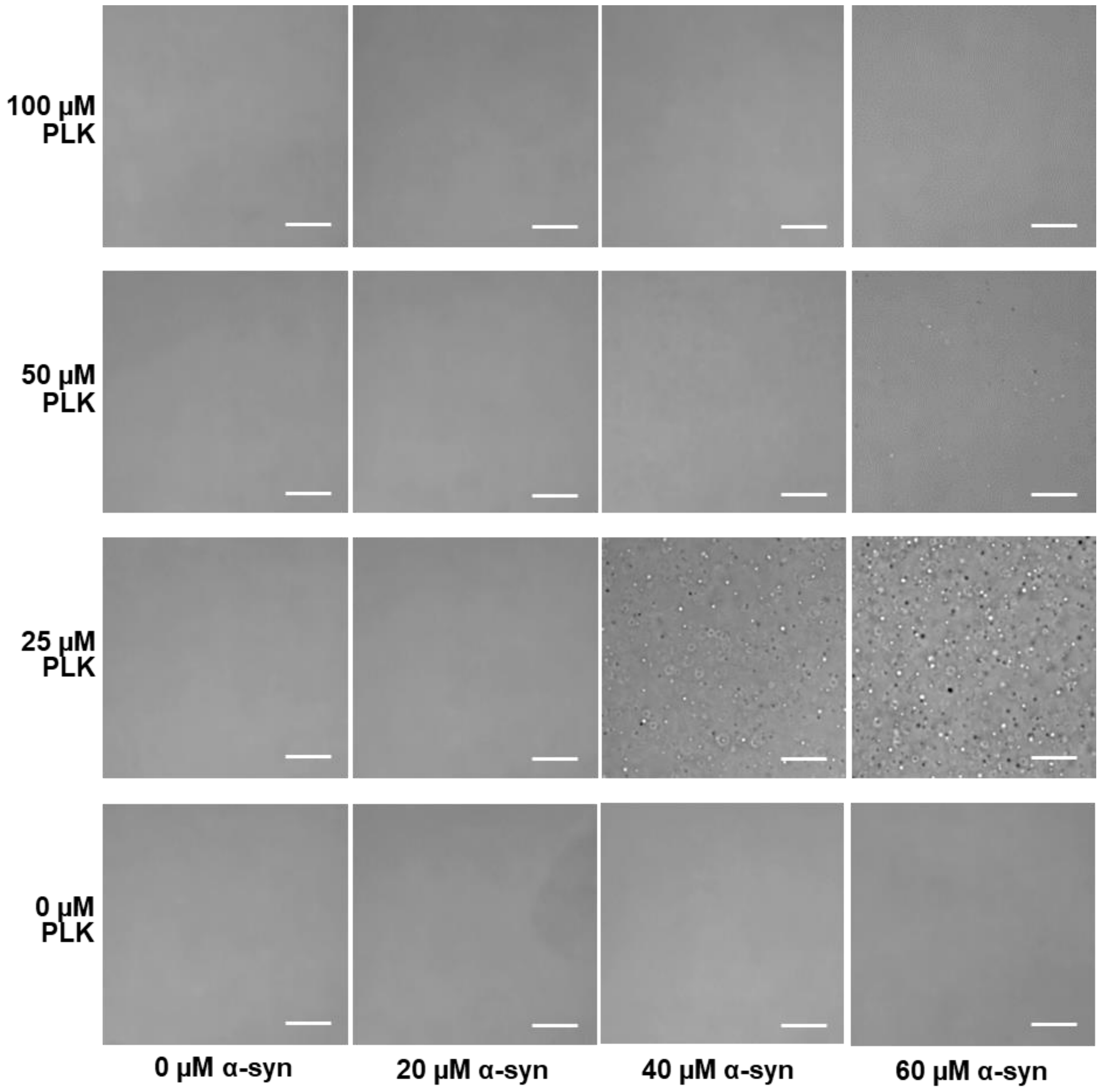


**Figure S5. Condensate formation depends on the ratio of FL α-syn to PLK.**

Representative DIC images of varying concentrations of FL α-syn and PLK in LLPS buffer. Images were taken 15 mins from PLK addition at 37 °C and used to generate a phase diagram by estimating the number of condensates per image. Scale bars represent 25 µm.

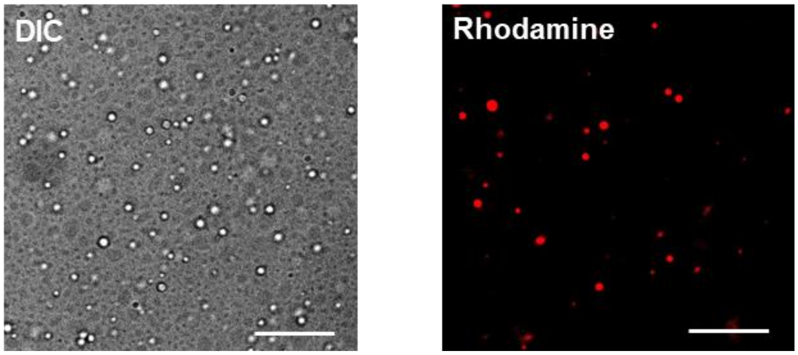


**Figure S6. α-Syn localizes within condensates.**

DIC (left) and Rhodamine fluorescence (right) images of 60 µM FL α-syn incubated at 37 °C with 25 µM PLK and 1 % rhodamine labelled FL α-syn in LLPS buffer. Images were taken 30 mins after PLK addition. Scale bars represent 20 µm.

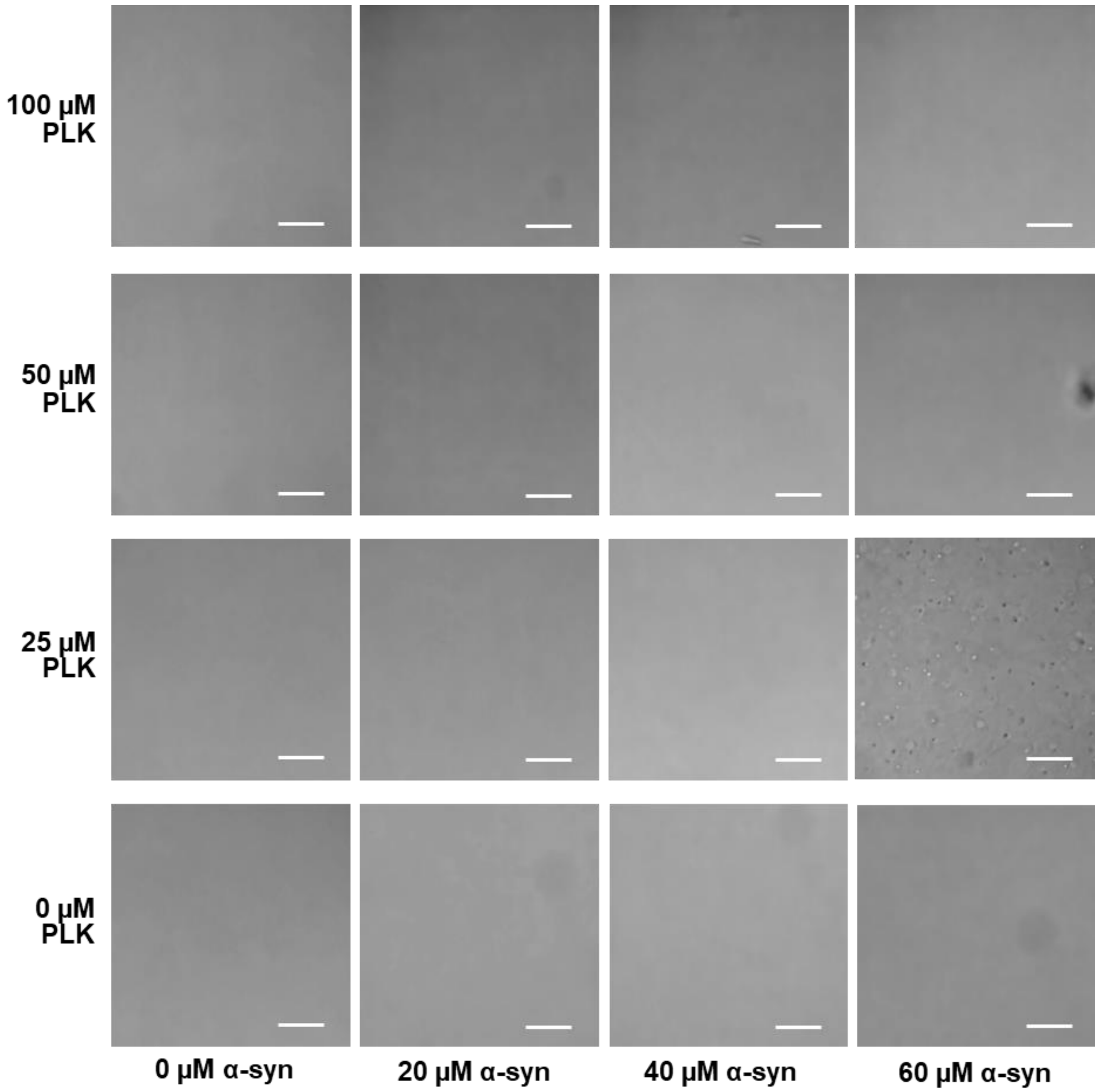


**Figure S7. 5-140 α-syn condensate formation depends on the ratio of protein to PLK.**

Representative DIC images of varying concentrations of 5-140 α-syn and PLK in LLPS buffer. Images were taken 15 mins from PLK addition at 37 °C and used to generate a phase diagram by estimating the number of condensates per image. Scale bars represent 25 μm.

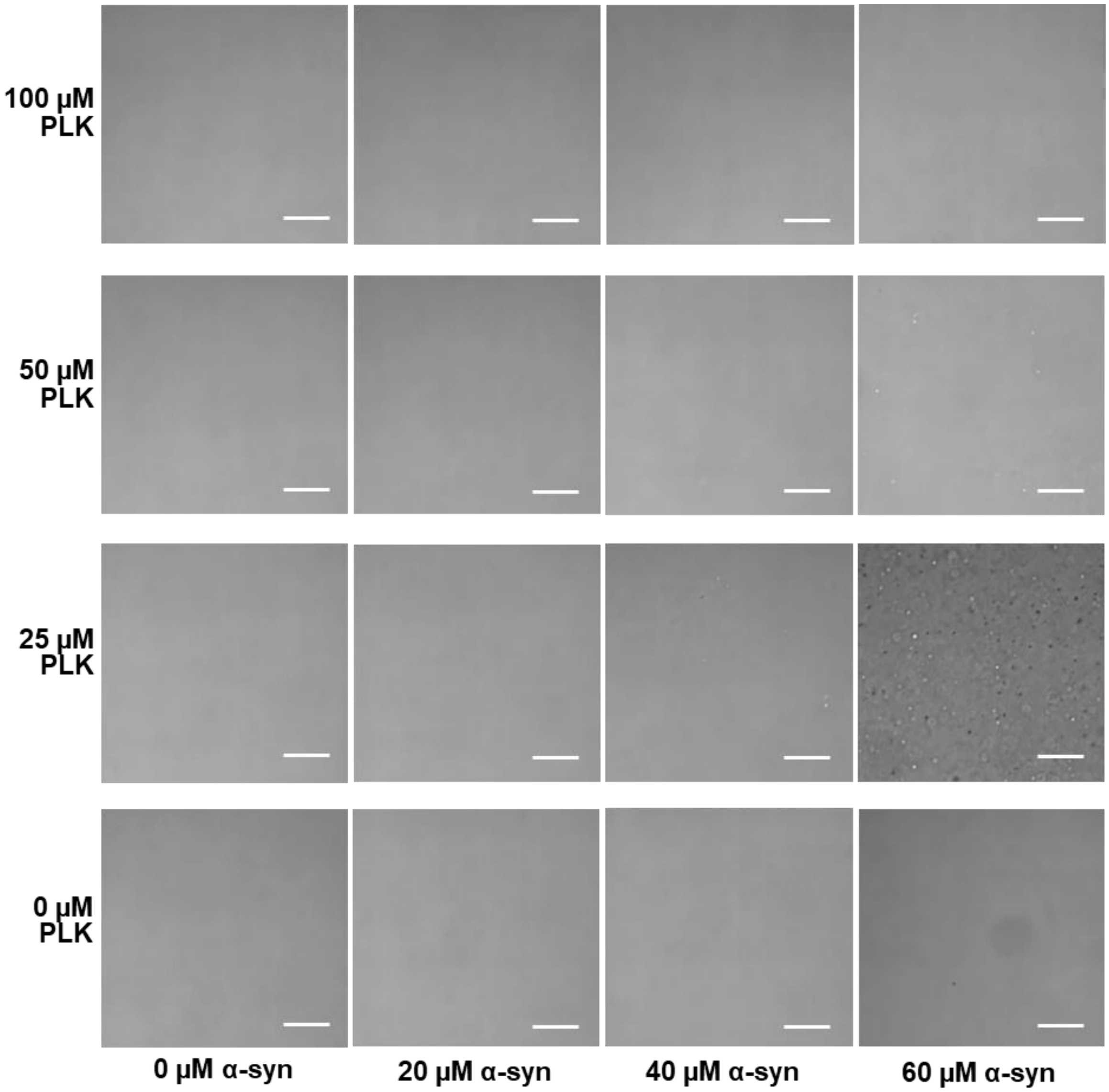


**Figure S8. 11-140 α-syn condensate formation depends on the ratio of protein to PLK.**

Representative DIC images of varying concentrations of 11-140 α-syn and PLK in LLPS buffer. Images were taken 15 mins from PLK addition at 37 °C and used to generate a phase diagram by estimating the number of condensates per image. Scale bars represent 25 µm.

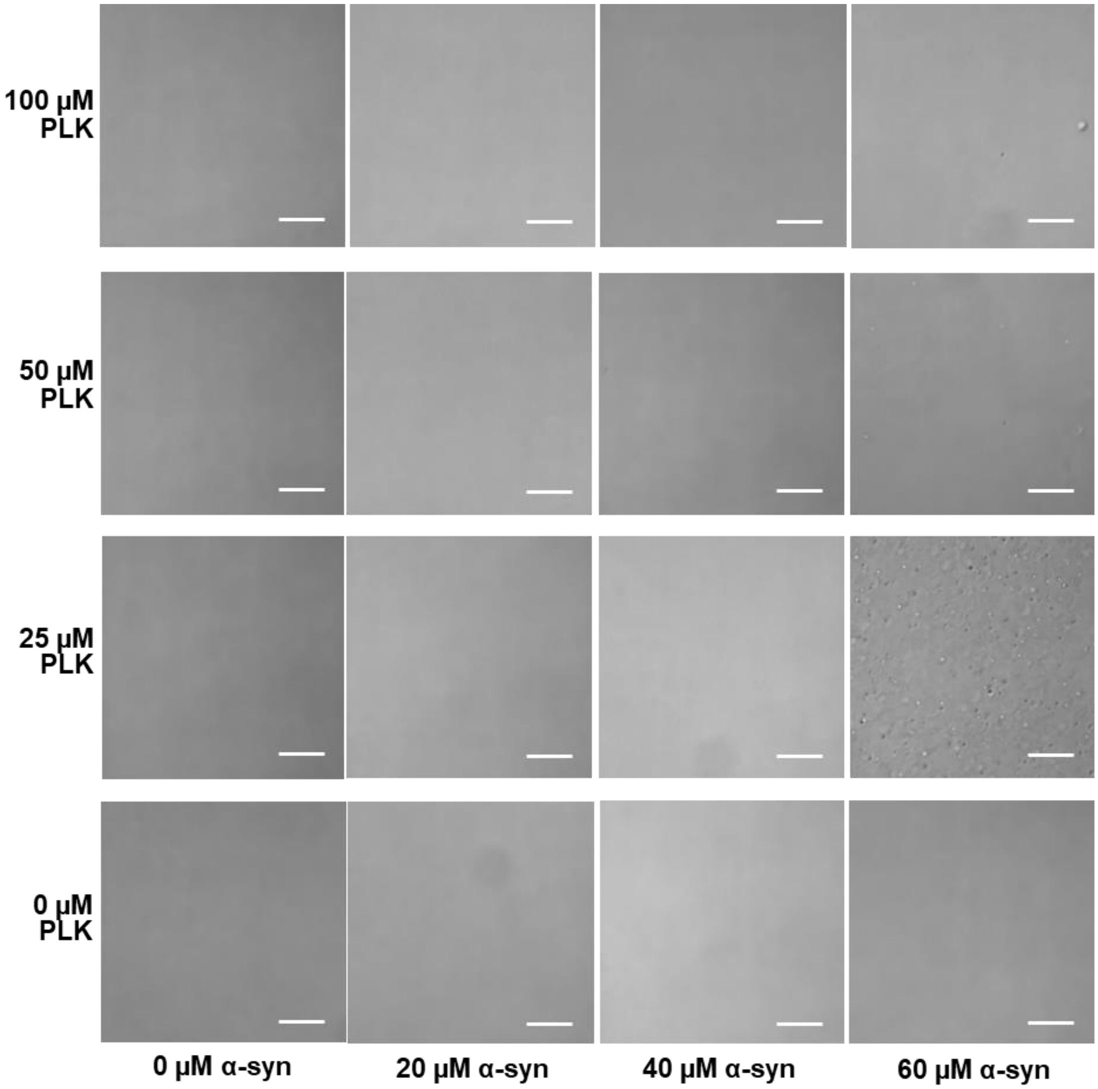


**Figure S9. 19-140 α-syn condensate formation depends on the ratio of protein to PLK.**

Representative DIC images of varying concentrations of 19-140 α-syn and PLK in LLPS buffer. Images were taken 15 mins from PLK addition at 37 °C and used to generate a phase diagram by estimating the number of condensates per image. Scale bars represent 25 μm.

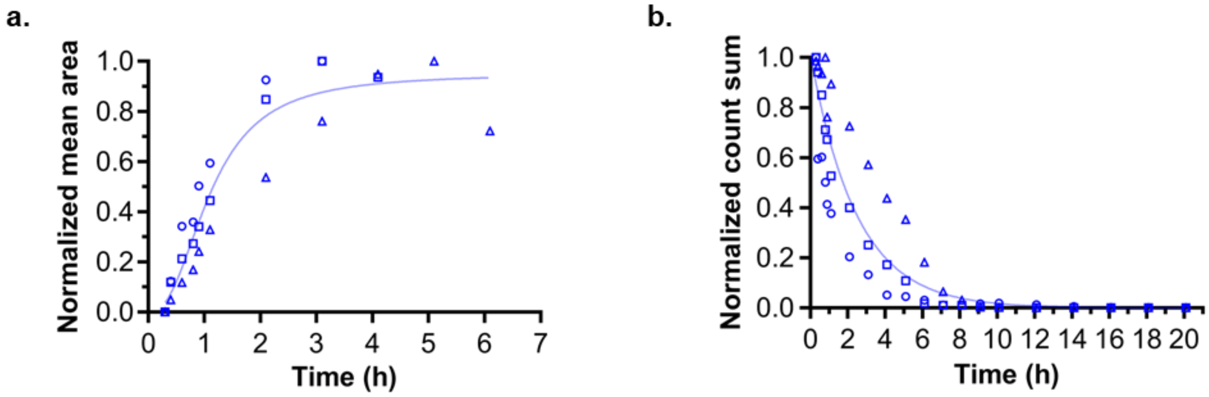


**Figure S10. FL α-syn condensates grown and reduce in number with time.**

a/b, Normalized (a) mean object area and (b) total object count against time for 60 µM FL α-syn incubated at 37 °C with 25 µM PLK in LLPS buffer. Three individual biological repeats are shown, represented by circular, square or triangular symbols. Data are globally fitted with (a) a one-phase association curve or (b) a one-phase decay curve (solid lines). Mean object area values for each time point are the normalized mean of all individual object area values compiled over the three z-stack images acquired. Time points with less than 10 % of the maximum number of objects were removed to reduce error resulting from small sample size. Total object count values for each time point are the normalized sum of the individual object counts measured in each of the three z-stack images acquired.

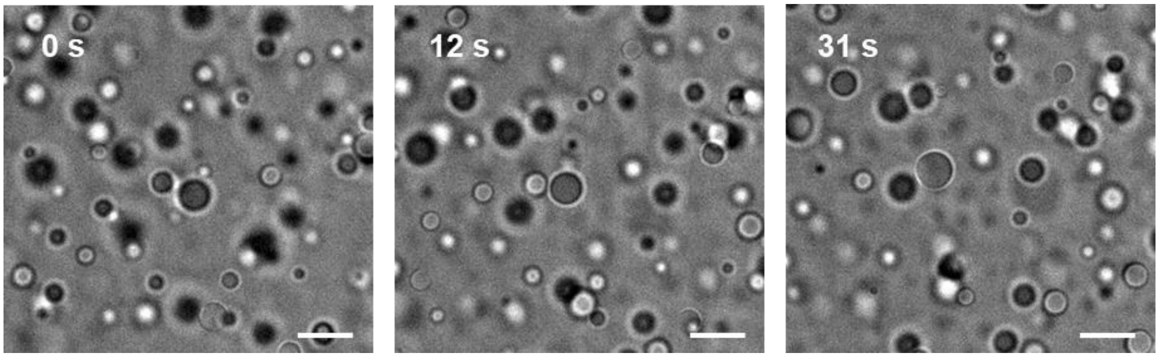


**Figure S11. Condensates can grow via coalescence.**

DIC images depicting a coalescence event of two condensates in solution for 60 µM FL α-syn with 25 µM PLK. Images were taken ~ 20 mins after PLK addition using a 60x oil objective. Scale bars represent 10 µm.

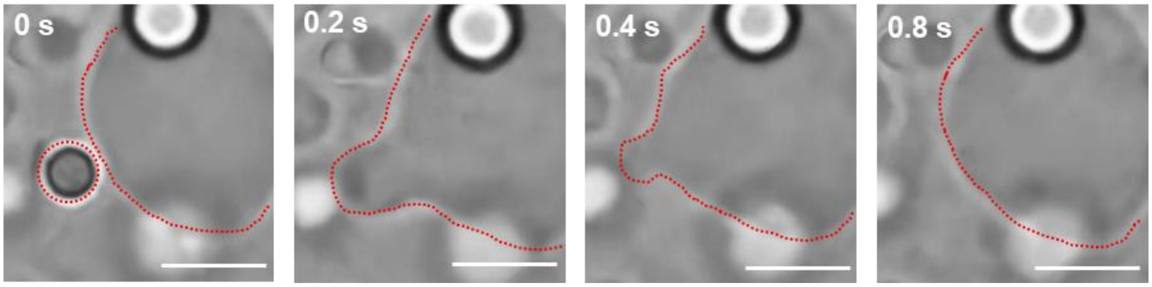


**Figure S12. Surface-wetted condensates can grow via coalescence.**

DIC images manually acquired at the bottom of the well after ~ 1.5 h incubation of 60 µM FL α-syn with 25 µM PLK. Images depict a coalescence event (scale bars represent 10 µm).

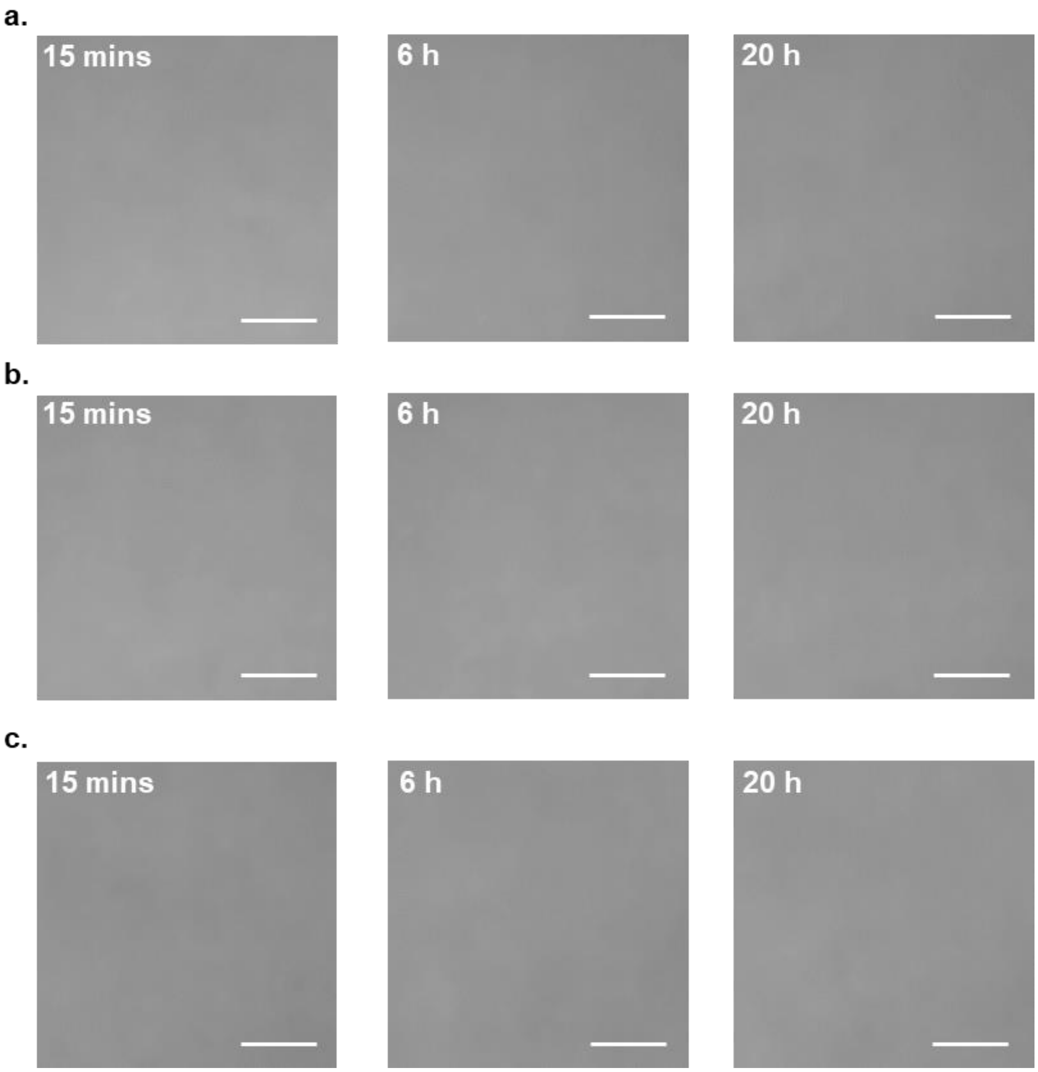


**Figure S13. Control samples do not form soluble condensates or aggregates.**

Representative DIC images of (a) LLPS buffer, (b) 25 µM PLK and (c) 60 µM FL α-syn at selected time points. Scale bars represent 25 µm.

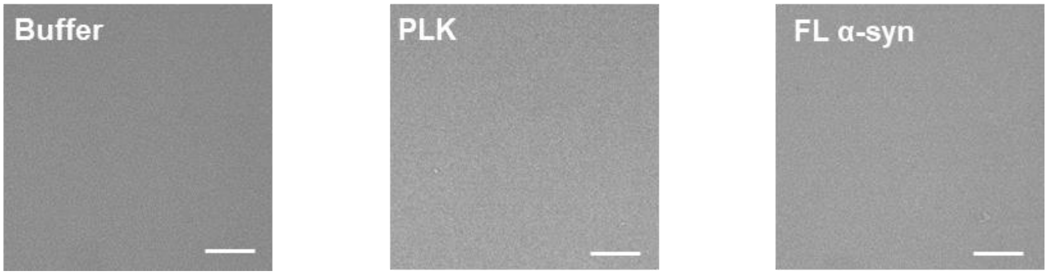


**Figure S14. Control samples do not form sedimented condensates or aggregates.**

Representative DIC images taken at the bottom surface of the well after 20 h incubation at 37 °C in LLPS buffer. Control samples of buffer, 25 µM PLK or 60 µM FL α-syn are shown. Scale bars represent 25 µm.

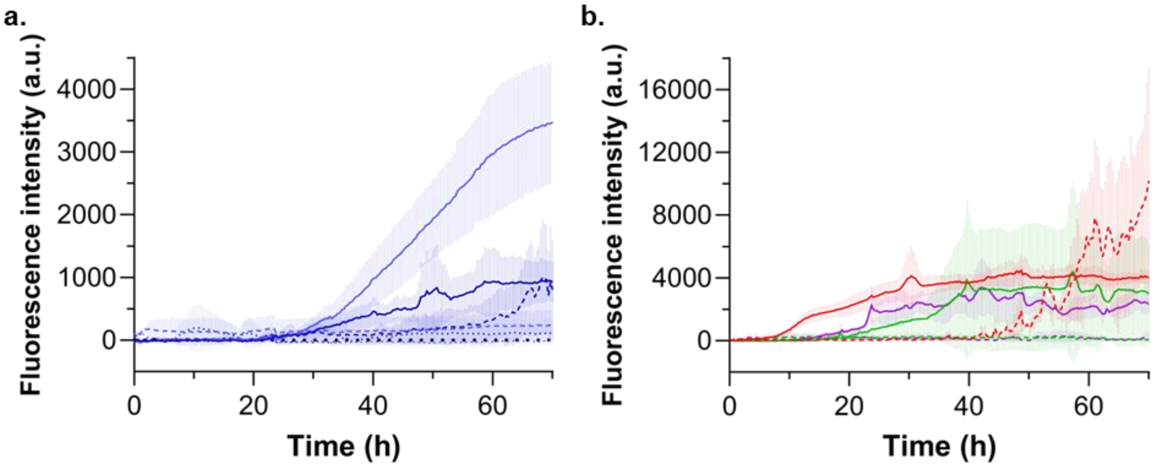


**Figure S15. N-terminal truncation accelerates aggregation following α-syn LLPS.**

a, Raw ThT fluorescence intensity data when 60 µM FL α-syn with 25 µM PLK (solid line), 60 µM FL α-syn (dashed line) and 25 µM PLK (dotted line), are incubated at 37 °C in LLPS buffer. Two independent biological repeats are shown in two shadows of blue. Each repeat is the mean of three technical replicates. Semi-transparent error bars represent the standard deviation of the mean. b, Raw ThT fluorescence intensity data when 5-140 (red), 11-140 (green) or 19-140 (purple) α-syn are incubated in LLPS buffer at 37 °C. 60 µM α-syn with 25 µM PLK (solid line) and 60 µM α-syn alone (dashed line) are shown. A single biological repeat is shown per protein variant. Each repeat is the mean of three technical replicates and semi-transparent error bars represent the standard deviation of the mean.

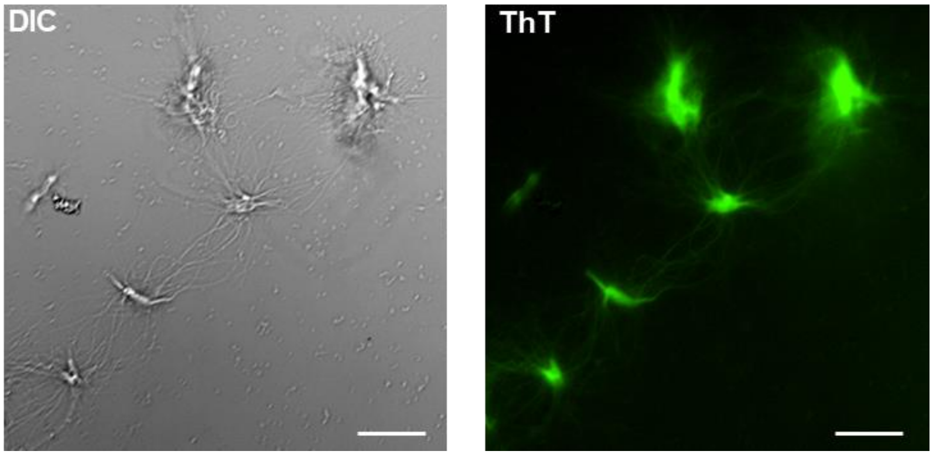


**Figure S16. N-terminal truncation accelerates aggregation following α-syn LLPS.**

Representative DIC (left) and fluorescent (right) images of 60 μM FL α-syn with 25 μM PLK at the endpoint of a LLPS ThT aggregation assay. Images are of the bottom of the well, scale bars represent 25 μm.

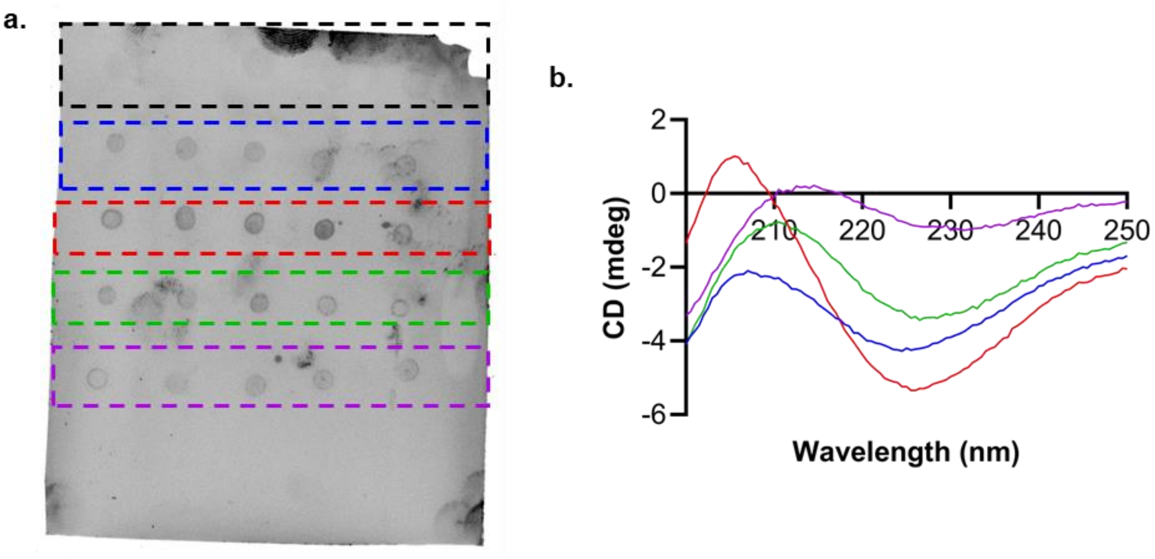


**Figure S17. Aggregates formed under LLPS conditions contain α-syn and β-sheet secondary structure**.

a, Representative dot-blot analysis of the insoluble aggregate pellet formed when FL (blue), 5-140 (red), 11-140 (green) and 19-140 (purple) α-syn were aggregated under LLPS conditions. As a control, analysis was also performed on a sample of 25 µM PLK in PBS (black), no fluorescence signal was observed for this sample confirming the specificity of the anti-α-syn primary antibody to α-syn. 5 replicates are shown per sample. b, Far-UV CD spectra of the insoluble aggregate pellet formed when FL (blue), 5-140 (red), 11-140 (green) and 19-140 (purple) α-syn were aggregated under LLPS conditions. The mean of five accumulations is shown per sample. As the aggregates formed under LLPS conditions may contain unknown concentrations of PLK, alongside α-syn, it was not possible to convert to MRE and so the raw data are shown here.

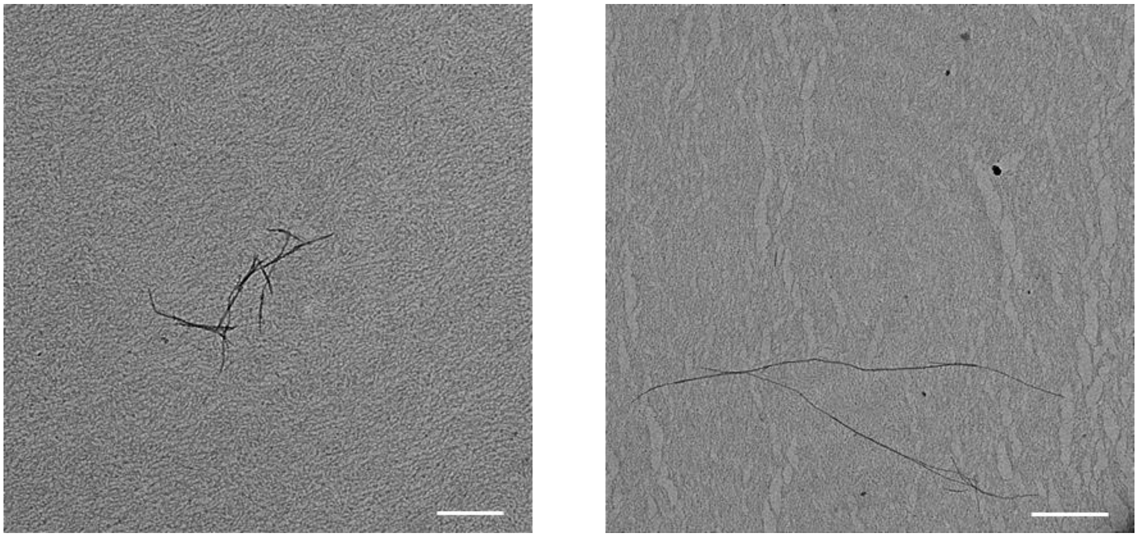

**Figure S18. α-syn aggregation is delayed in the absence of LLPS.**

Representative TEM images of 60 µM FL α-syn aggregated alone (i.e., in the absence of PLK) in LLPS buffer supplemented with 20 µM ThT. An aliquot of the whole sample was taken at the endpoint of a LLPS ThT aggregation assay and applied to a TEM grid. Scale bars represent 500 nm (left) and 1 µm (right).

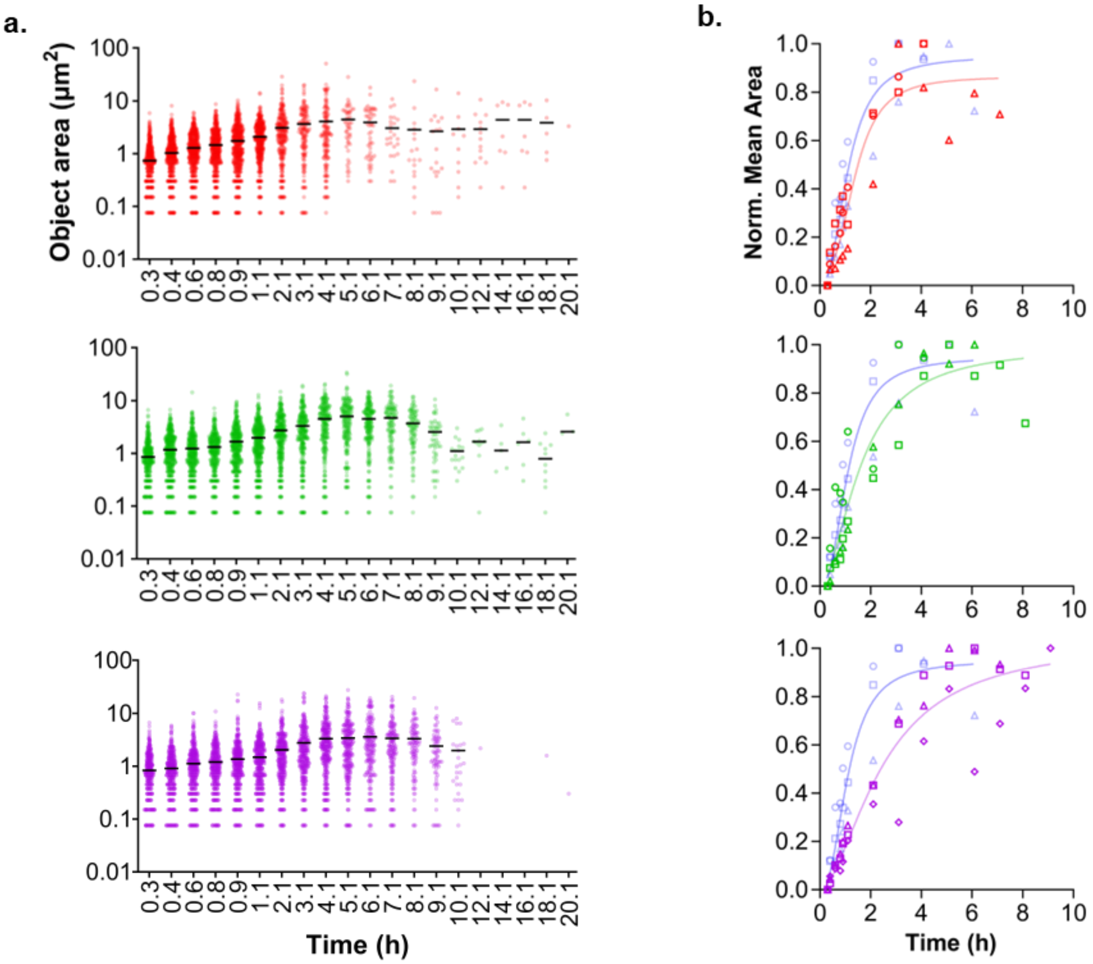


**Figure S19. N-terminal truncation increases α-syn condensate lifetime in solution.**

a, Object area distribution over time for 5-140 (red), 11-140 (green) and 19-140 (purple) α-syn. Each timepoint was compiled over the three z-stack images. Mean droplet areas at each time point are indicated by a solid black line. b, Normalized mean object area against time for 60 μM 5-140 (red), 11-140 (green) and 19-140 (purple) α-syn incubated with 25 μM PLK in LLPS buffer. Three individual biological repeats are shown, represented by circular, square or triangular symbols. The data is globally fit with a one-phase association curve (solid line). Mean object area values for each time point are the normalized mean of all individual object area values compiled over the three z-stack images acquired. Time points with less than 10 % of the maximum number of objects have been removed to reduce error resulting from smaller sample sizes. Each plot is overlayed with the corresponding FL α-syn data, shown in semi-transparent blue, for comparison.

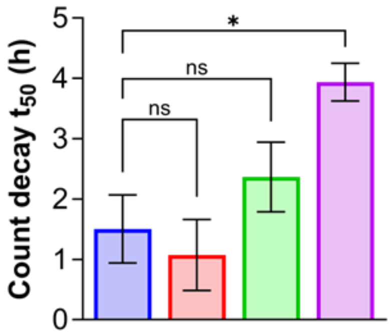


**Figure S20. Increasing N-terminal truncation increases α-syn condensate lifetime in solution.**

Half-life (t50) of the decay in object count for 60 µM FL (blue), 5-140 (red), 11-140 (green) and 19-140 (purple) α-syn with 25 µM PLK in LLPS buffer. Error bars represent the standard error of the mean for three biological repeats.

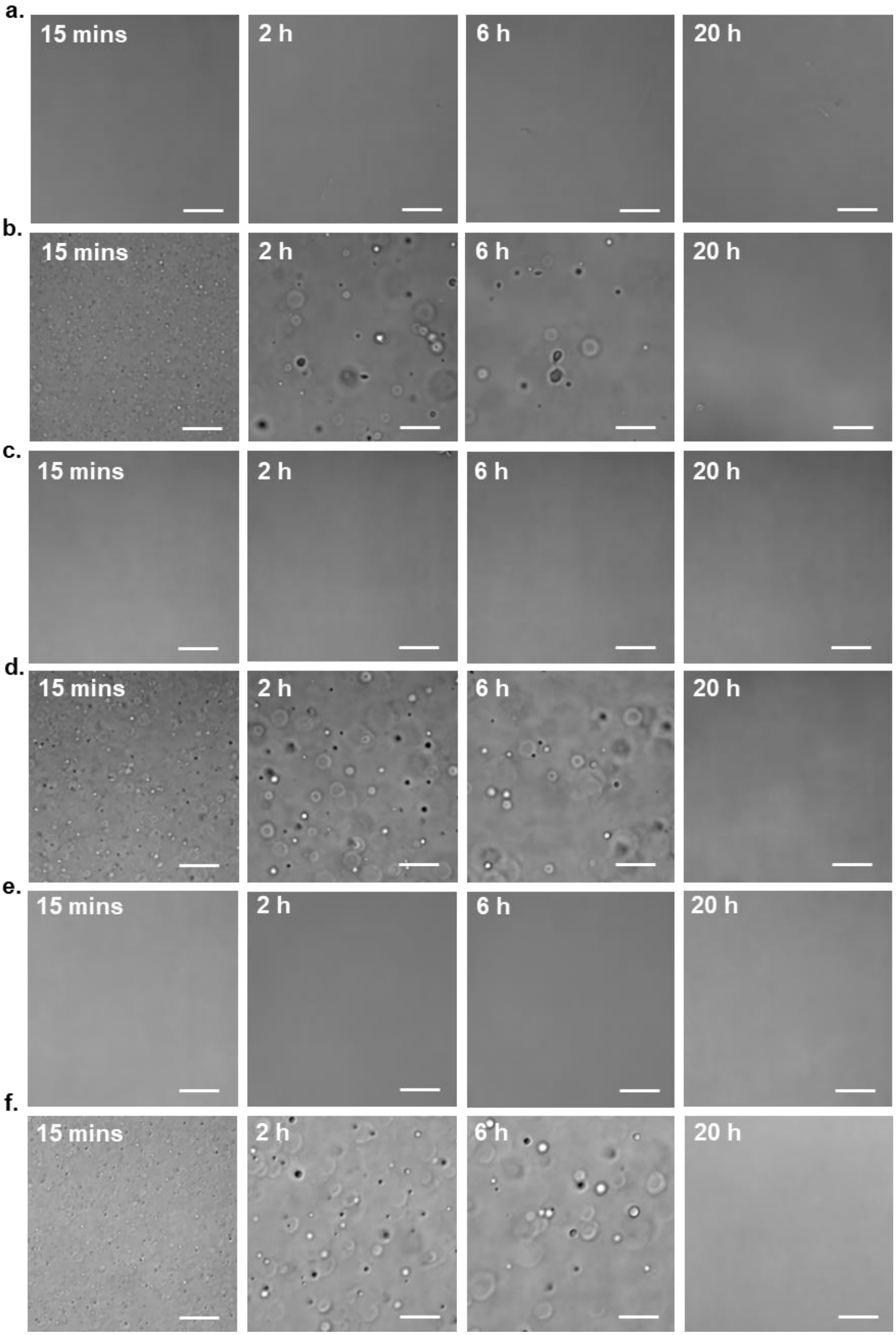


**Figure S21. N-terminally truncated α-syn condensates grow and decrease in number over time.**

Representative DIC images of (a) 60 µM 5-140 α-syn, (b) 60 µM 5-140 α-syn with 25 µM PLK, (c) 60 µM 11-140 α-syn, (d) 60 µM 11-140 α-syn with 25 µM PLK, (e) 60 µM 19-140 α-syn, and (f) 60 µM 19-140 α-syn with 25 µM PLK. Selected time points are shown. No condensate or aggregate formation was observed in the bulk solution in the absence of PLK for all variants. Scale bars represent 25 µm.

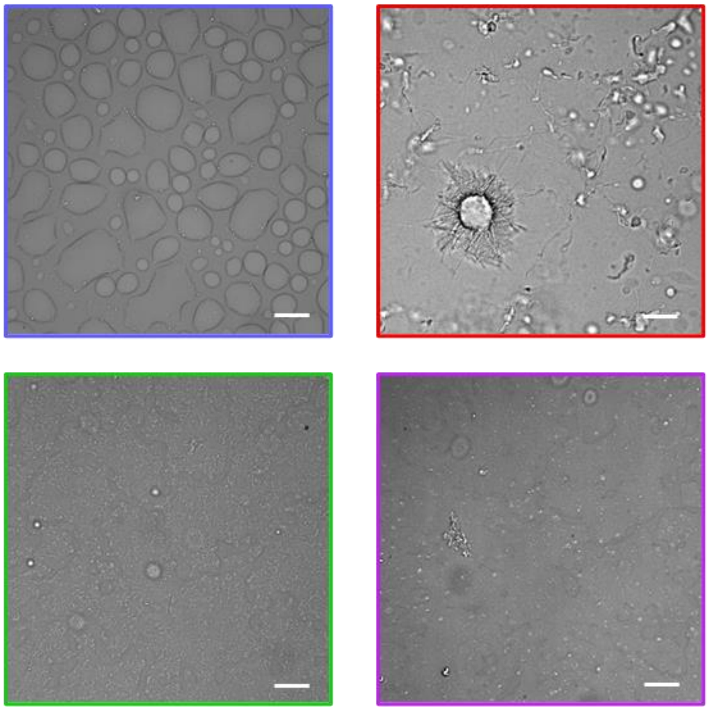

**Figure S22. N-terminally truncated α-syn condensates grow and decrease in number over time.**

Representative DIC images acquired manually at the bottom of the well after 20 h incubation of FL (blue), 5-140 (red), 11-140 (green) and 19-140 (purple) α-syn under LLPS conditions (scale bars represent 25 μm).

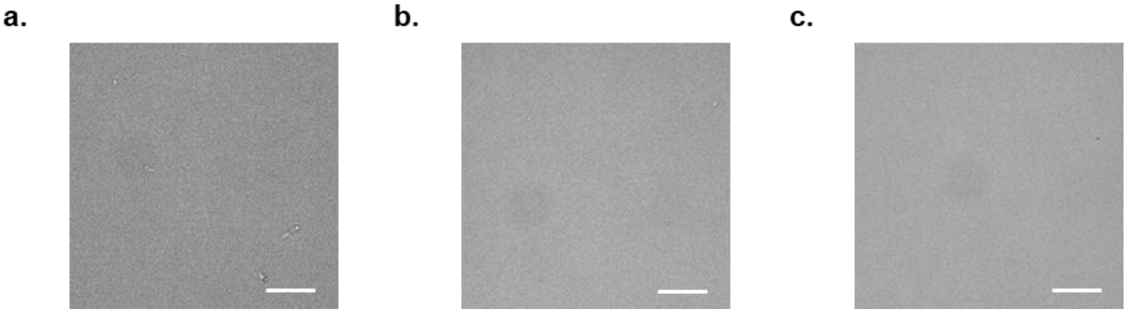

**Figure S23. N-terminally truncated α-syn alone does not form sedimented condensates or aggregates.**

Representative DIC images taken at the bottom of the well after 20 h incubation of 60 µM (a) 5-140, (b) 11-140 and (c) 19-140 α-syn at 37 °C in LLPS buffer. Scale bars represent 25 µm.

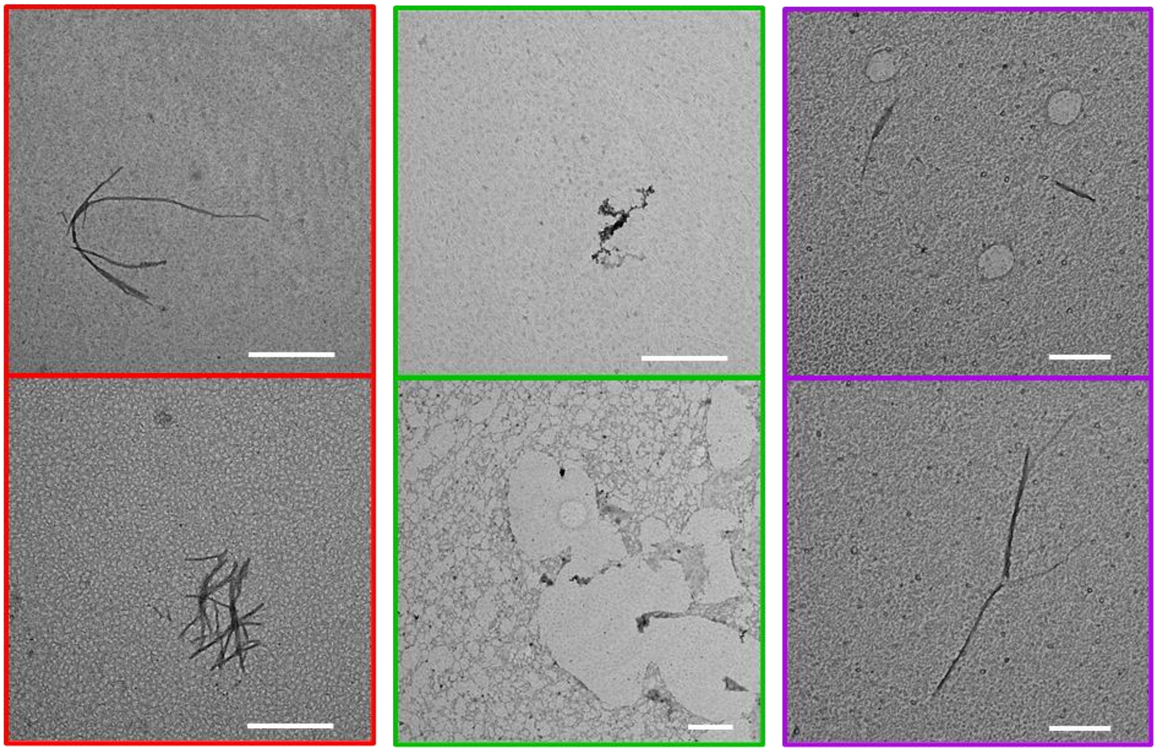

**Figure S24. N-terminally truncated α-syn aggregation is delayed in the absence of LLPS.**

Representative TEM images of 60 µM 5-140 (red), 11-140 (green) and 19-140 (purple) α-syn aggregated alone (i.e., in the absence of PLK) in LLPS buffer with 20 µM ThT. An aliquot of the whole sample was taken at the endpoint of a LLPS ThT aggregation assay and applied to the TEM grid. Scale bars represent 500 nm.

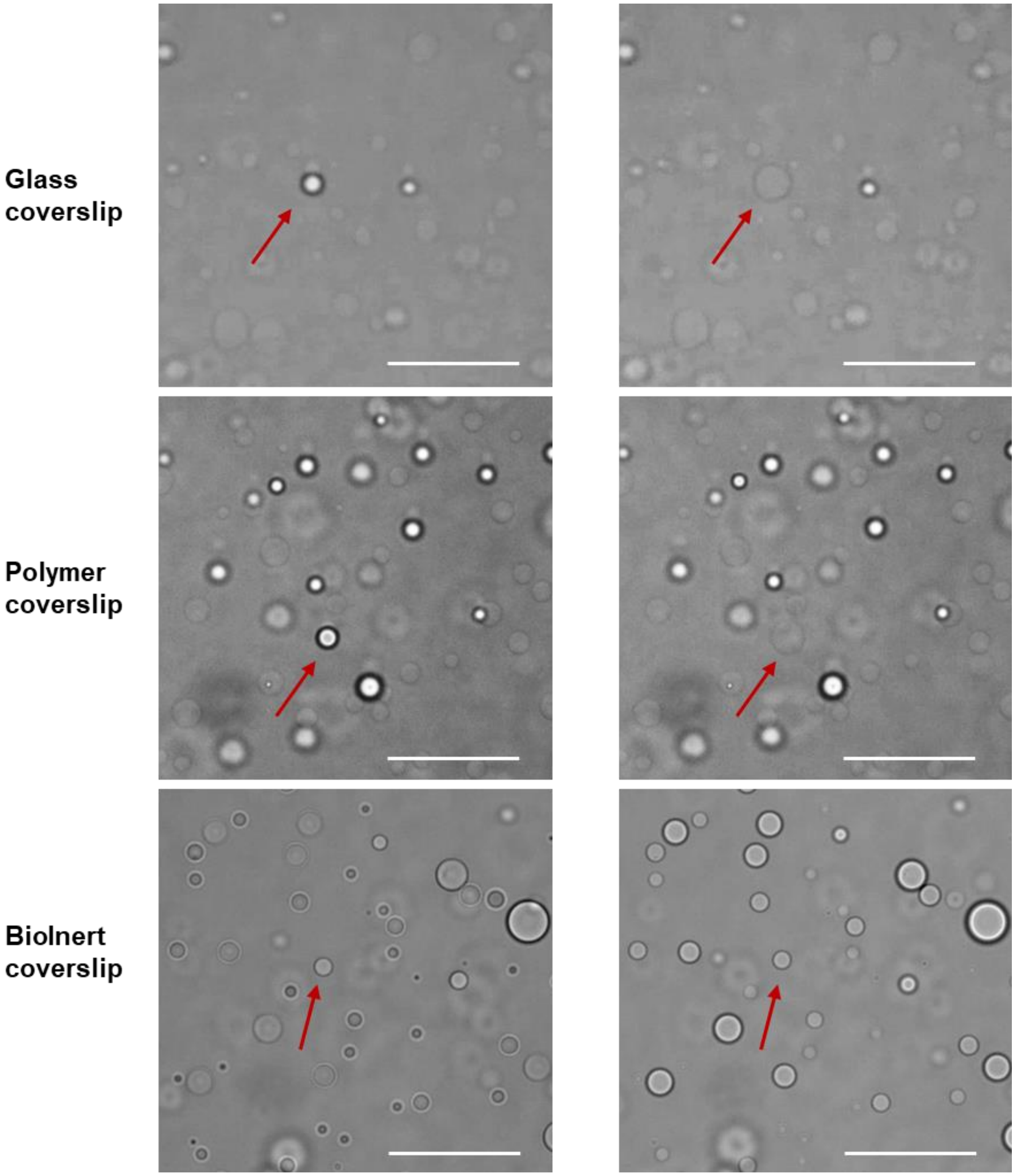


**Figure S25. Increased surface hydrophobicity decreases the wettability of α-syn condensates.**

Representative DIC images of 60 µM FL α-syn with 25 µM PLK in LLPS buffer incubated at 37 °C in 8-well slides different coverslips. Images were acquired at the bottom surface of the well before (left) and after (right) a condensate in solution (indicated by a red arrow) wet the well surface. Scale bars represent 25 µm.

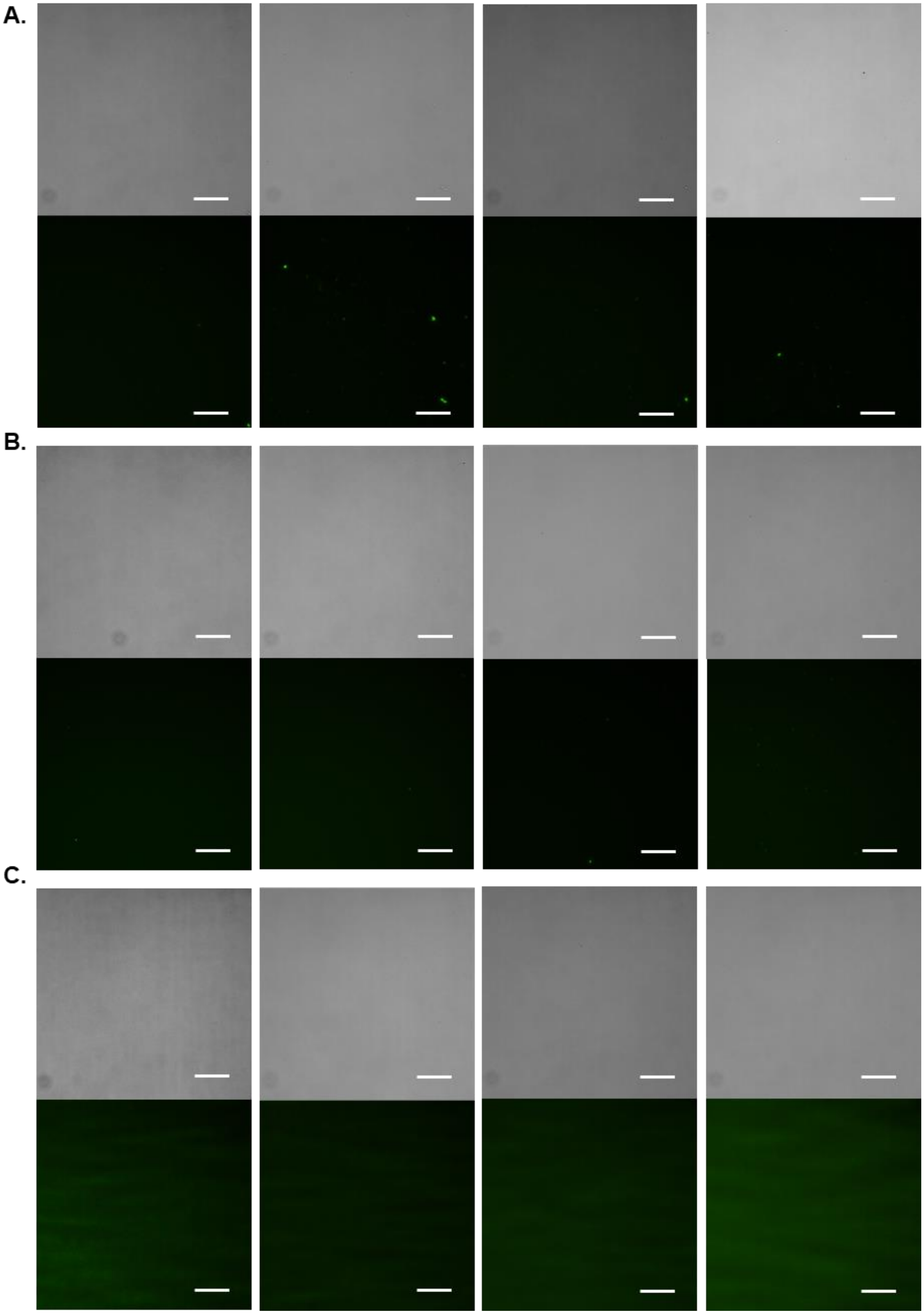

**Figure S26. Control samples for the surface hydrophobicity screening assay.**

(A,B,C) Representative DIC (top) and fluorescent (bottom) images of LLPS buffer incubated at 37 °C in 8-well slides with a glass (A), polymer (B) or BioInert (C) coverslip. Images were acquired at selected time points at the bottom surface of the well. Scale bars represent 25 µm.

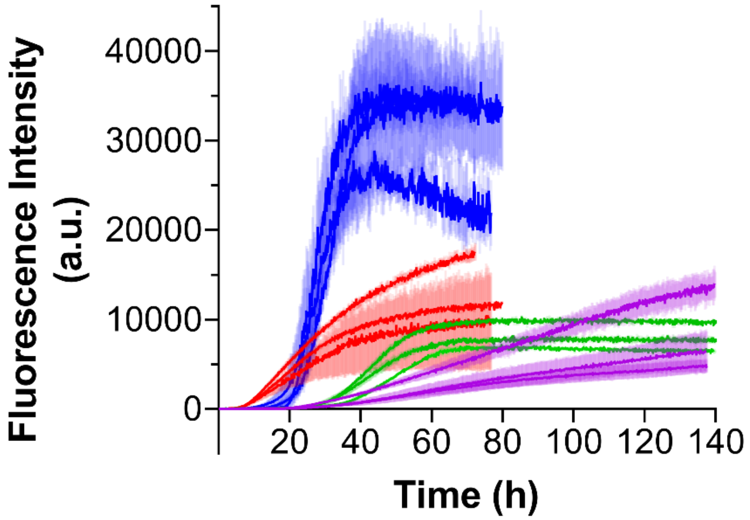


**Figure S27. N-terminal truncation modulates α-syn aggregation.**

Raw dispersed solution amyloid aggregation of FL (blue), 5-140 (red), 11-140 (green) and 19-140 (purple) α-syn, monitored by ThT fluorescence intensity at 37 °C with agitation. Three individual biological repeats are shown per variant. Each repeat is the mean of either two or three technical replicates and semi-transparent error bars represent the standard deviation of the mean.

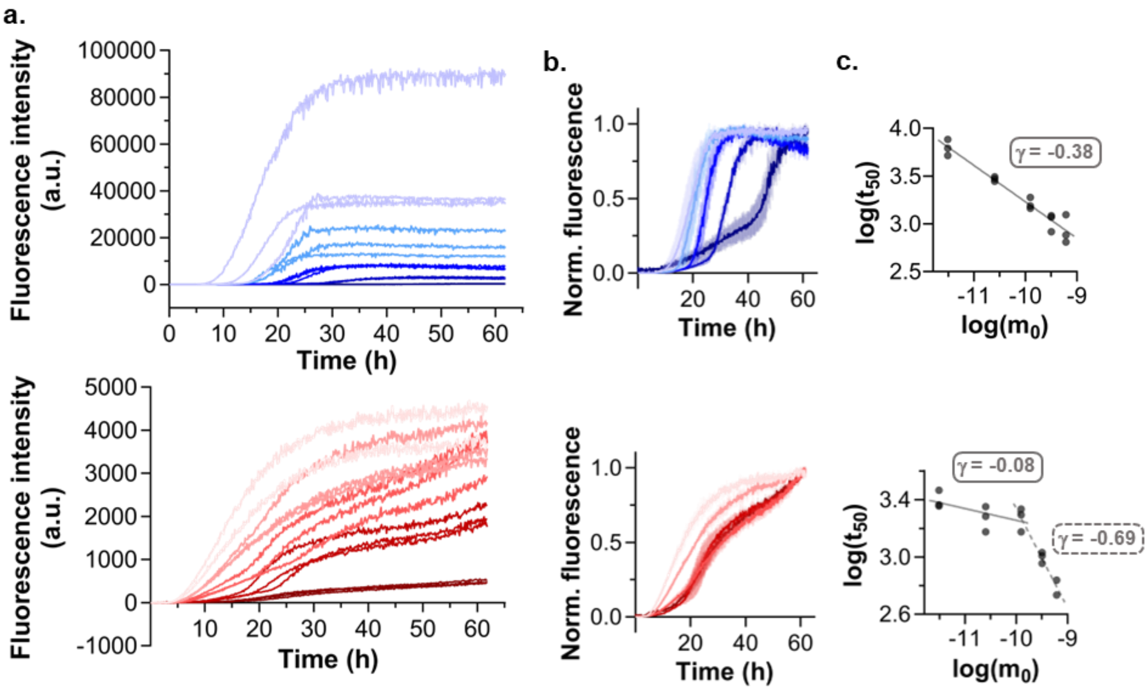


**Figure S28. 5-140 α-syn aggregation has an increased secondary nucleation contribution.**

a, Raw dispersed solution amyloid aggregation data for varied concentrations (10, 25, 50, 75 and 100 µM) of FL (top, dark blue to light blue) and 5-140 (bottom, dark red to light red) α-syn, monitored by ThT fluorescence intensity at 37 °C with agitation. n ≥ 2 technical replicates from a single assay are shown. b, Normalization of the data shown in (a) for FL (top) and 5-140 (bottom) α-syn, where each curve is the mean of the technical replicates and semi-transparent error bars represent the standard deviation, n ≥ 2. c, Plot of the log of initial monomer concentration (m0) versus the log of the half-life of aggregation (t50), used to estimate the scaling exponent (γ) and provide insight into the mechanism of aggregation. n ≥ 2 technical replicates from a single assay are shown, corresponding to FL (top) and 5-140 (bottom) α-syn.

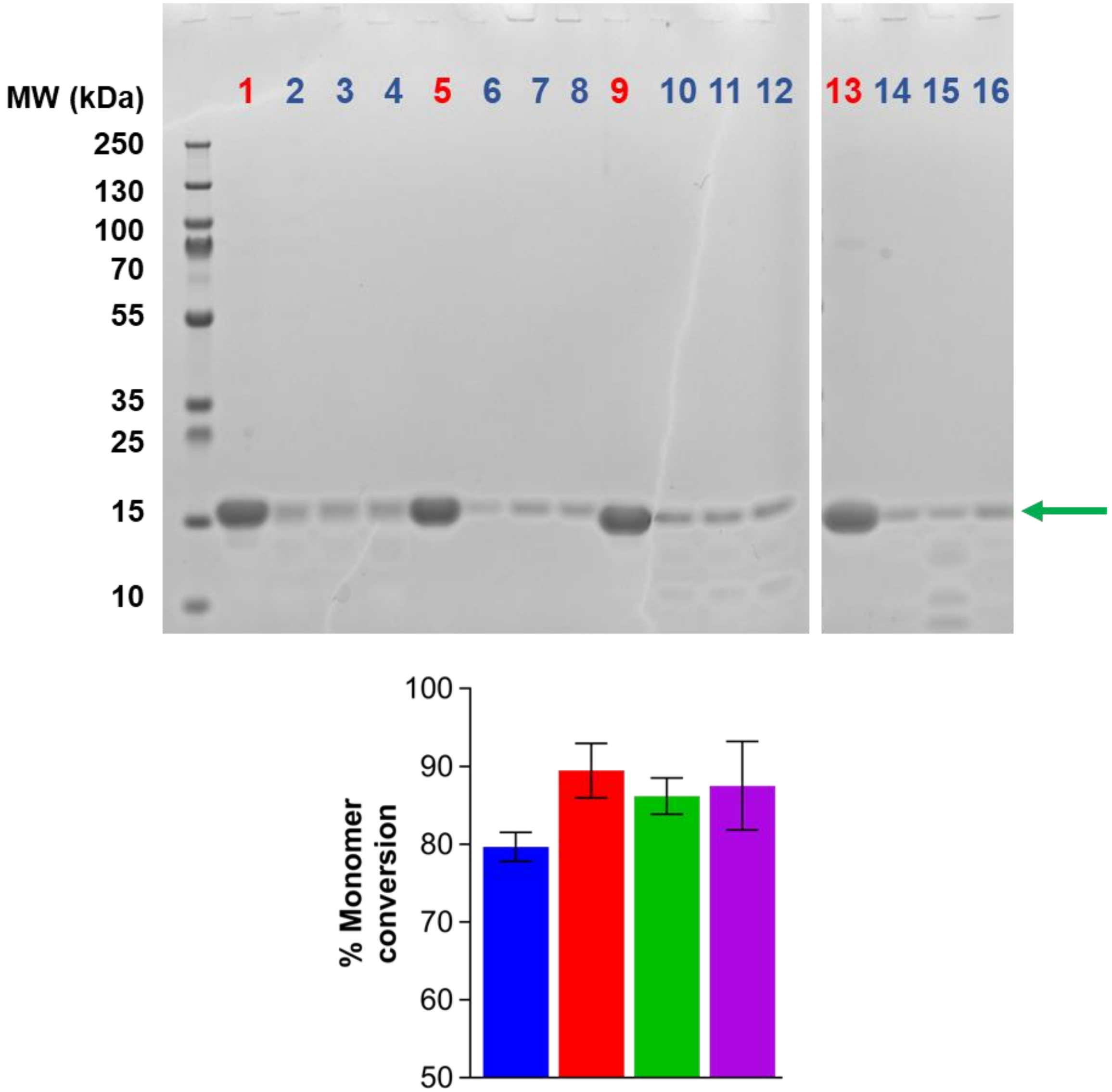


**Figure S29. N-terminal truncation does not affect α-syn monomer conversion.**

Top: Representative SDS PAGE image of the soluble fraction of FL (lane 1-4), 5-140 (lane 5-8), 11-140 (lanes 9-12) and 19-140 (lanes 13-16) α-syn at the start (red) and end (blue) of a dispersed solution aggregation assay. Green arrow indicates the band quantified. The band intensity, background corrected using a blank area of gel, was measured using Fiji to estimate % conversion of α-syn monomer to insoluble aggregate. Bottom: corresponding quantification of the percentage conversion of soluble FL (blue), 5-140 (red), 11-140 (green) and 19-140 (purple) α-syn into insoluble aggregates during a dispersed solution aggregation assay. Representative biological repeat is shown. Each repeat is the mean of three technical replicates and semi-transparent error bars represent the standard deviation of the mean.

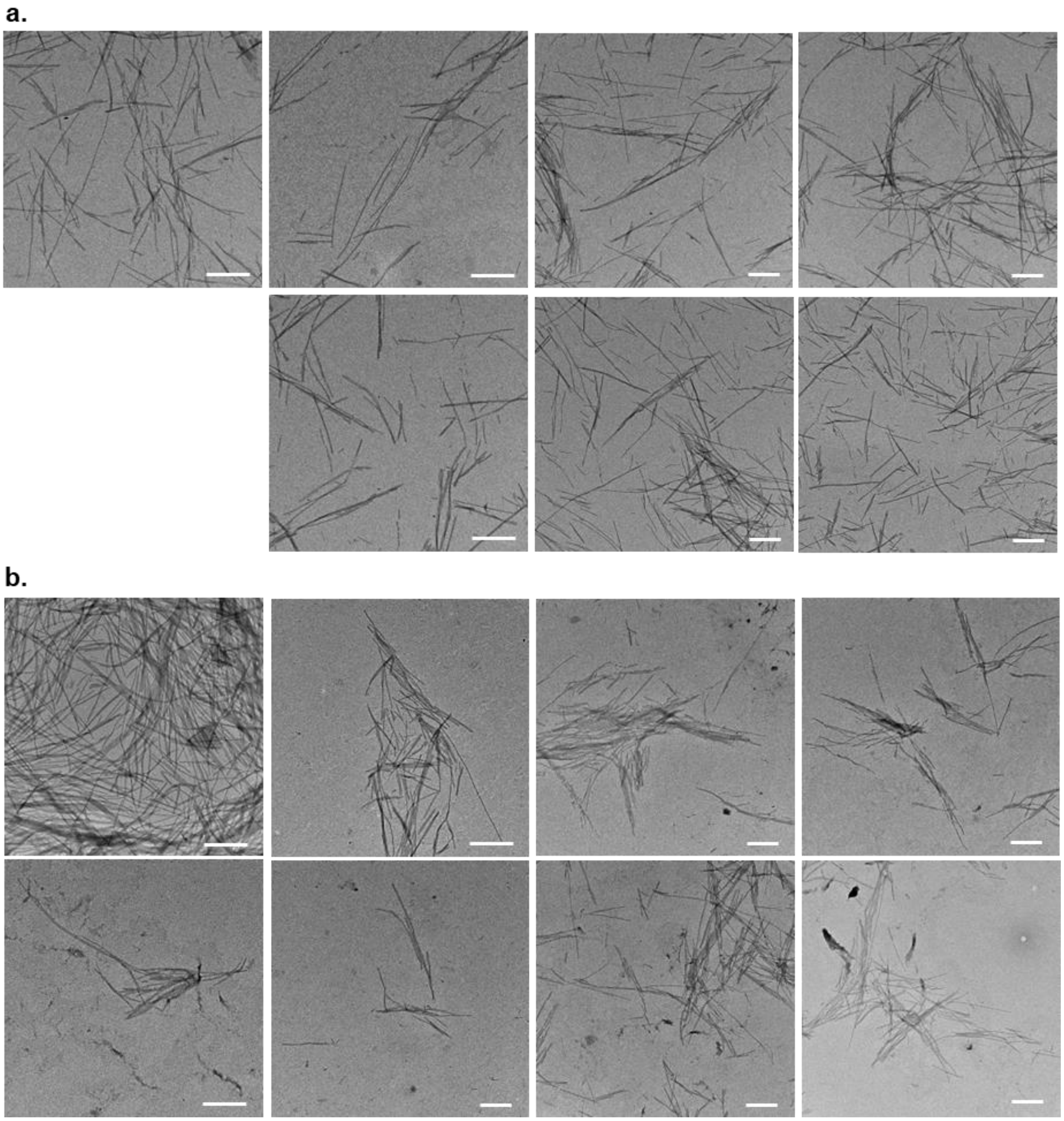
a.
b.

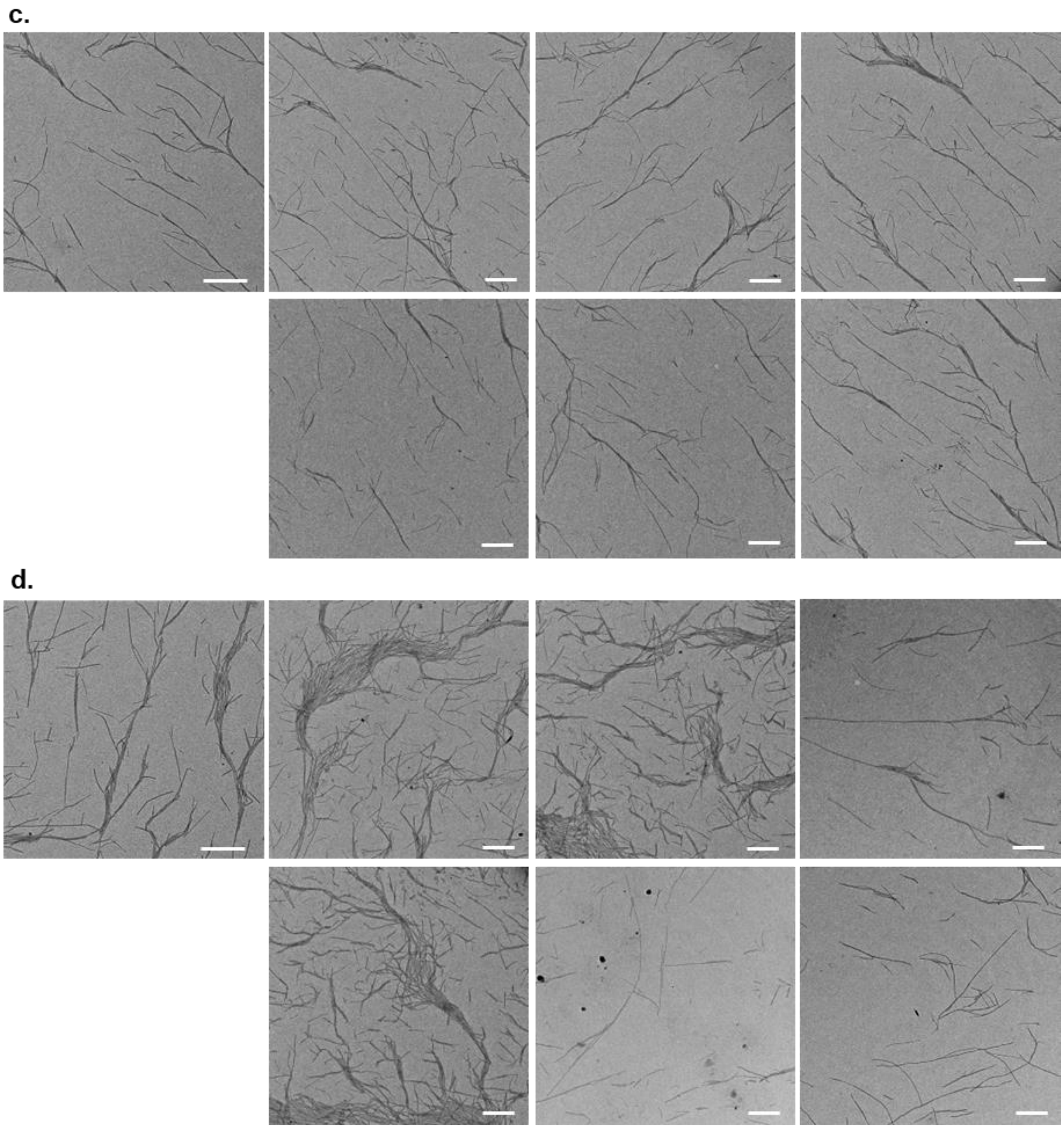


**Figure S30. N-terminal truncation affects α-syn amyloid morphology.**

TEM images of FL (a), 5-140 (b), 11-40 (c) and 19-140 (d) α-syn taken at the end point of the dispersed solution aggregation assay and used for quantitative analysis of fibril length and width by Fiji. Scale bars represent 500 nm.

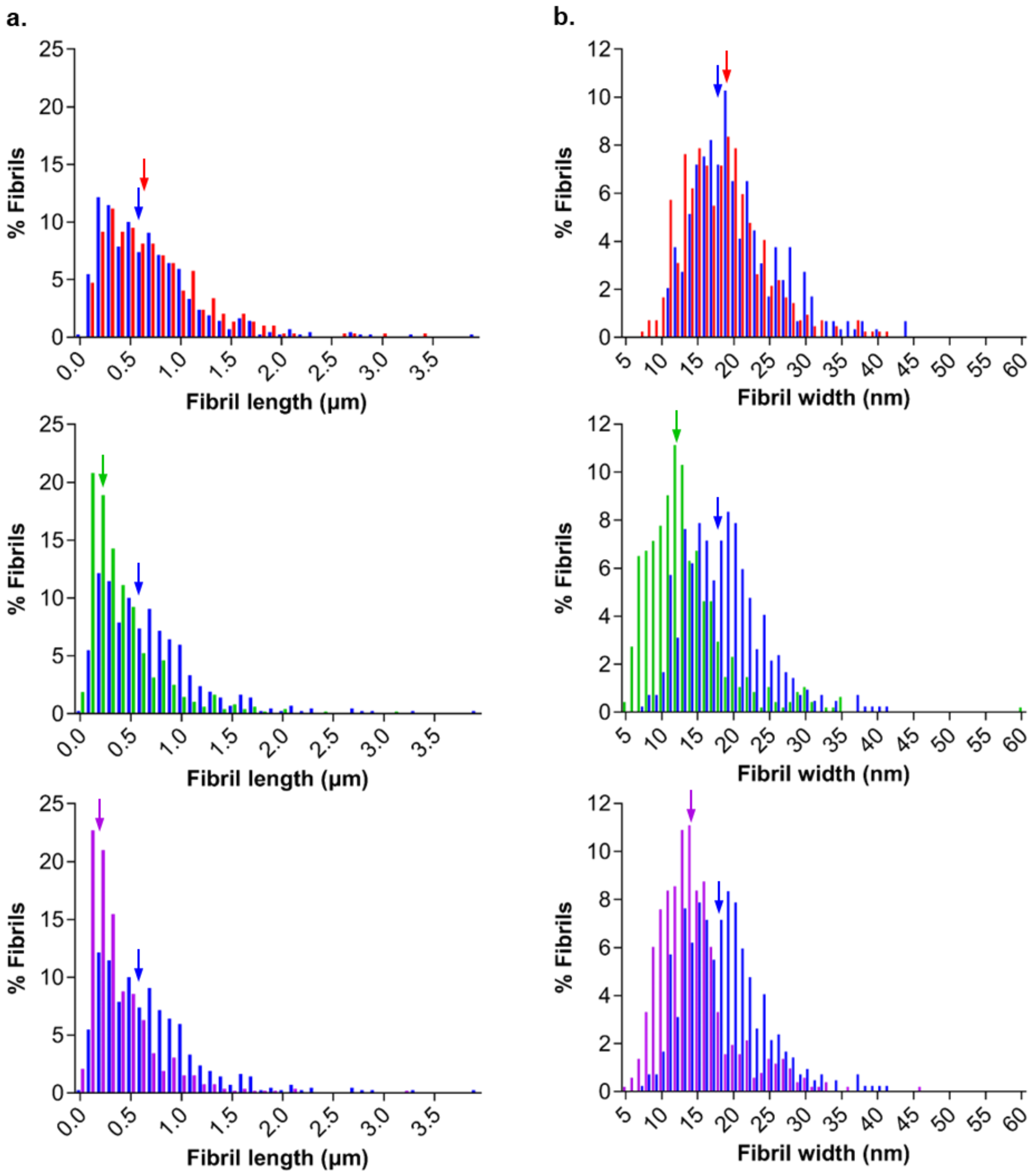


**Figure S31. Increased N-terminal truncation decreases the length and width of α-syn amyloids.**

Length (a) and width (b) frequency distributions of FL (blue), 5-140 (red), 11-140 (green) and 19-140 (purple) α-syn amyloids as determined by quantitative analysis performed on TEM images of the insoluble fraction at the endpoint of a dispersed solution aggregation assay (length: n = 419, 295, 476 and 524 respectively, width: n = 419, 292, 476 and 514 respectively). Colored arrows indicate the corresponding median value for each variant. All TEM images quantified are shown in Figure 4 or Figure S26.

FL and 5-140 α-syn fibrils are similar in length (median lengths 0.59 and 0.63 µm respectively), whereas 11-140 and 19-140 α-syn aggregates are typically shorter (median lengths 0.31 and 0.28 µm, respectively). 5-140 and FL α-syn do not display a significant difference in the frequency distributions of their fibril thickness, with medians of 18 and 19 nm, respectively. In contrast, 11-140 and 19-140 α-syn fibrils are thinner (median widths 12 and 14 nm, respectively).

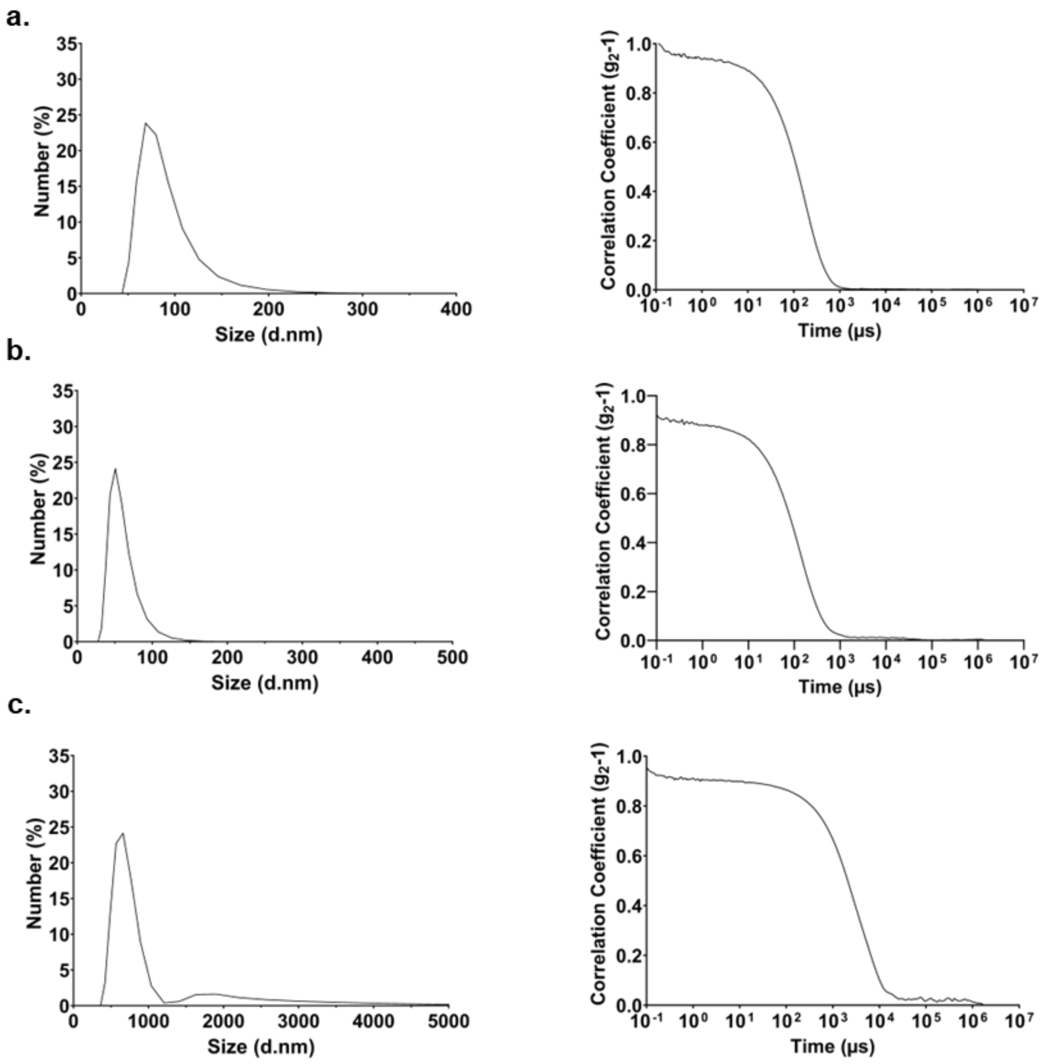


**Figure S32. Lipid vesicle and pre-formed fibril size confirmed by DLS.**

Representative DLS data of (a) the DMPS lipid vesicles used to study primary nucleation, (b) short FL α-syn fibrils used for fibril elongation, or (c) long FL α-syn fibrils used for secondary nucleation. Number distribution is shown on the left and the corresponding correlogram on the right.
