## Supplementary Movie Legends for "Surface Wetting Is a Key Determinant of α-Synuclein Condensate Maturation"

### Legends for Supplementary Movies 1 to 4

**Supplementary Movie 1 (separate file). FL α-syn aggregate formation following LLPS.**

z-stack movie taken at the endpoint of a ThT assay under LLPS conditions when 60 µM FL α-syn was incubated with 25 µM PLK in LLPS buffer at 37 °C. The movie is an overlay of both DIC and ThT fluorescence channels and shows the formation of a large fibril network that extends across the bottom of the well and into solution. 2.5 µm z step, 64 µm z range, scale bar represents 50 µm.

**Supplementary Movie 2 (separate file). 5-140 α-syn aggregate formation following LLPS.**

z-stack movie taken at the endpoint of a ThT assay under LLPS conditions when 60 µM 5-140 α-syn was incubated with 25 µM PLK in LLPS buffer at 37 °C. The movie is an overlay of both DIC and ThT fluorescence channels and shows the formation of a large fibril network that extends across the bottom of the well and into solution. 2.5 µm z step, 64 µm z range, scale bar represents 50 µm.

**Supplementary Movie 3 (separate file). 11-140 α-syn aggregate formation following LLPS.**

z-stack movie taken at the endpoint of a ThT assay under LLPS conditions when 60 µM 11-140 α-syn was incubated with 25 µM PLK in LLPS buffer at 37 °C. The movie is an overlay of both DIC and ThT fluorescence channels and shows the formation of a large fibril network that extends across the bottom of the well and into solution. 2.5 µm z step, 64 µm z range, scale bar represents 50 µm.

**Supplementary Movie 4 (separate file). 19-140 α-syn aggregate formation following LLPS.**

z-stack movie taken at the endpoint of a ThT assay under LLPS conditions when 60 µM 19-140 α-syn was incubated with 25 µM PLK in LLPS buffer at 37 °C. The movie is an overlay of both DIC and ThT fluorescence channels and shows the formation of a large fibril network that extends across the bottom of the well and into solution. 2.5 µm z step, 64 µm z range, scale bar represents 50 µm.
